# Small molecules exploiting structural differences within microRNA-200 precursors family members reverse a type 2 diabetes phenotype

**DOI:** 10.1101/2020.06.27.175281

**Authors:** Hafeez S. Haniff, Xiaohui Liu, Laurent Knerr, Malin Lemurell, Daniel Abegg, Alexander Adibekian, Matthew D. Disney

**Affiliations:** The Scripps Research Institute, Department of Chemistry, 130 Scripps Way, Jupiter, FL 33458, USA; Medicinal Chemistry, Research and Early Development Cardiovascular, Renal and Metabolism, BioPharmaceuticals R&D, AstraZeneca, Gothenburg, Pepparedsleden, 1, SE-431 83 Mölndal, Sweden

## Abstract

MicroRNA families are pervasive in the human transcriptome, but specific targeting of individual members is a challenge because of sequence homology. Many of the secondary structures of the precursors to these miRs (pre-miRs), however, are quite different. Here, we demonstrate both *in vitro* and *in cellulis* that design of structure-specific small molecules can inhibit specific miR family members to modulate a disease pathway. In particular, the miR-200 family consists five miRs, miR-200a, −200b, −200c, −141, and - 429, and is associated with Type II Diabetes (T2D). We designed a small molecule that potently and selectively targets pre-miR-200c’s structure. The compound reverses a pro-apoptotic effect in a pancreatic β-cell model. In contrast, oligonucleotides targeting the RNA’s sequence inhibit all family members. Global proteomics analysis further demonstrates selectivity for miR-200c. Collectively, these studies establish that miR-200c plays an important role in T2D and that small molecules targeting RNA structure can be an important complement to oligonucleotides targeting sequence.

**Significance Statement:** The most common way to develop medicines targeting RNA is by using oligonucleotides that target its sequence by using base pairing. Some RNAs, however, have similar sequences and thus are impossible to target selectively by using oligonucleotides. Here, we show that a class of RNAs that have similar sequences emerge from precursors that have very different structures. Exploiting these structural differences afforded a selective compound. In particular, the selective small molecule targets a member of the microRNA (miR)-200 family, the overexpression of which is linked to diabetes and pancreatic cell death. Selective inhibition of family member miR-200c alleviates pancreatic cell death, and thus the small molecule provides a path to the treatment of diabetes.

---

RNA structure influences and controls biology, lending the biomolecule its diverse functions. The ability to target RNA structure with synthetic ligands has been a strategy to dissect and understand RNA biology and to leverage it for the development of therapeutics (1-4). One class of RNAs particularly amendable to targeting with synthetic ligands is non-coding microRNAs (miRs) (5). These small regulatory RNAs are synthesized as a primary microRNA transcripts (pri-miRs) that are processed in the nucleus by the microprocessor complex Drosha-DGCR8, affording precursor microRNAs (pre-miRs). The pre-miRs are translocated to the cytoplasm and processed by the nuclease Dicer liberating the mature microRNAs (miR) (6). The miRs bind to mRNAs with sequence complementarity in the 3’ untranslated region (3’ UTR), suppressing translation (7).

Three approaches have been used to develop miR inhibitors: (i) oligonucleotide-based antagomirs to bind to the mature miRs via base pairing (8); (ii) small molecules that target pri- or pre-miR structures found at functional Dicer or Drosha processing sites, thereby inhibiting biogenesis (9-11); and (iii) small molecules that target miR precursor structures to trigger cleavage of the RNA target (12-14). One of the challenges with targeting some miRs specifically with oligonucleotides is that family members can have the same or similar sequences (15, 16). The structures of the precursors to these miRs, however, can be quite different (15, 16). The identification of ligands that exploit these differences can thus provide a way to selectively inhibit one miR family member to study its biology and provide leads for precision medicines. Herein, we study the ability to differentially affect the biology of miR-200c, a member of the miR-200 family containing miR-200a, −200b, and −200c, −141, and −429.

Importantly, the miR-200 family contributes to type 2 diabetes (T2D) and, in particular, is causative of β-cell apoptosis (17, 18). Indeed, overexpression of miR-200c or miR-141 is sufficient to induce β-cell apoptosis *in vivo* (17, 18). Furthermore, β-cell apoptosis can be reduced by ablation of the entire miR-200 cluster (18). The miR-200c family is broken down into two subfamilies based on sequence similarity of the seed sequence that forms base pairing interactions with mRNA 3’ UTRs: miR-200b, −200c, and −429 form one sub-family while miR-141 and −200a form the other (19, 20). This overlap makes dissecting the contribution of each miR to β-cell -function and survival challenging.

Herein, we show that oligonucleotide-based approaches that target RNA sequence cannot discriminate broadly against the miR-200 family because of sequence homology. Analysis of the precursors from which the miRs emanate, however, shows broad differences in the RNA structures that could be exploited with structure-specific organic ligands (**Figure 1**). Indeed, we designed a structure-binding ligand that is a selective inhibitor of pre-miR-200c biogenesis. Further, the compound exerts selective and potent effects on the miRnome and the proteome and stimulates beneficial effects on a T2D-associated phenotype in a β-cell model. Thus, these studies validate that miR-200c is a phenotype-driving biomolecule.

**Figure 1:**
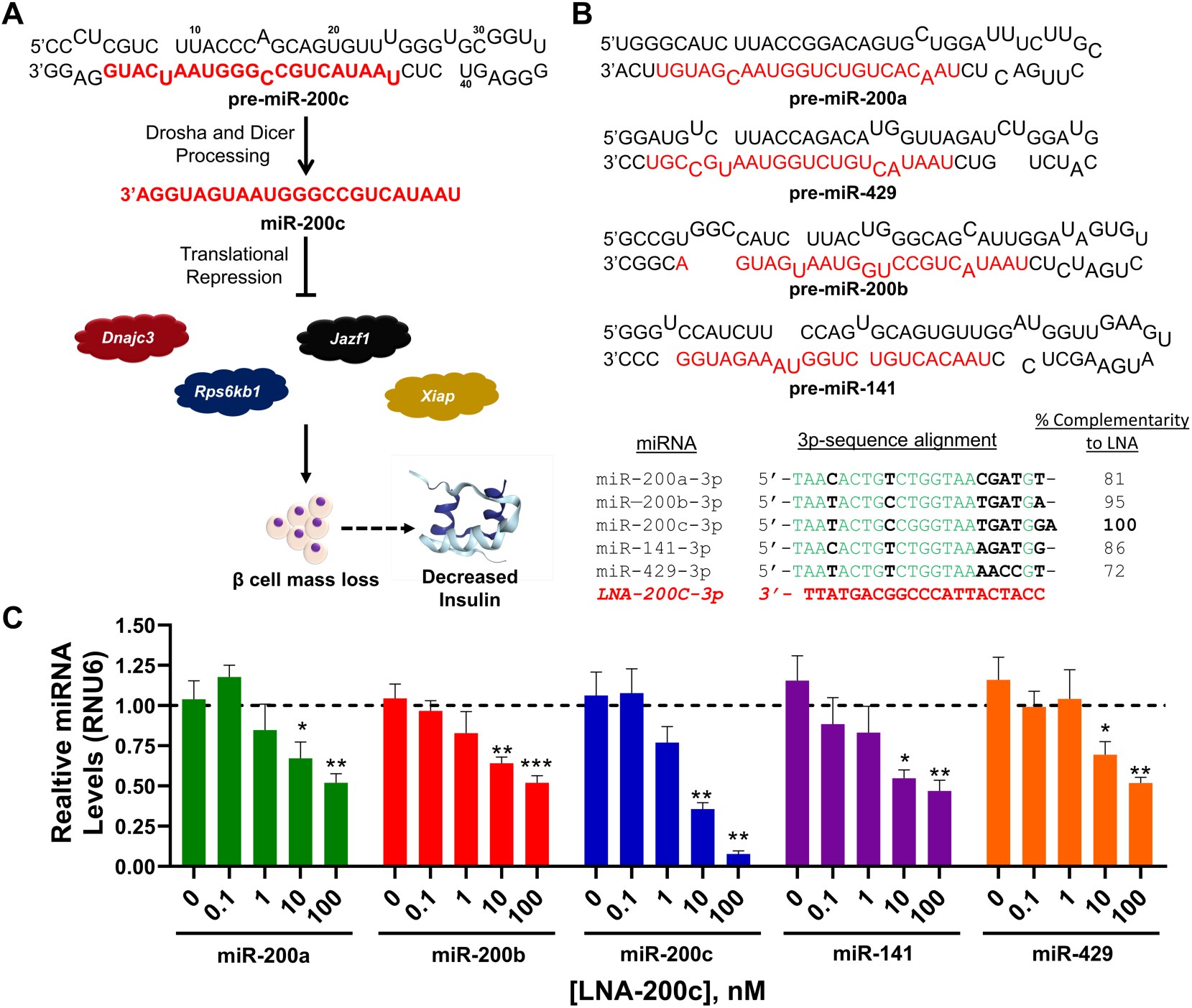
Schematic of the miR-200c pathway, sequence similarity of the miR-200c family, and effect of a miR-200c-targeting antagomir. **A**) Precursor miR-200c is processed by Dicer to generate mature miR-200c and translationally represses Dnajc3, Rps6kb1, Jazf1, and Xiap. These proteins regulate the activation of apoptosis and thus their down-regulation triggers apoptosis of β-cells and decreases insulin. **B**) The miR-200 family consists of miR-200a, miR-200b, miR-200c, miR-141 and miR-429 which each form structurally distinct hairpins but have extensive sequence homology in their mature sequences. **C**) Effect of LNA-200c on all miRNAs in the miR-200 family, which all miR-200 family members as they have >70% sequence complementarity to the LNA (n = 4). *, p<0.05; **, p<0.01. All p-values were calculated by a two-tailed Student t-test. All data are reported as mean ± S.E.M.

## Results

### Analysis of the miRNA-200 family to design structure-specific ligands

MiRNA families are a class of non-coding RNAs that have similar functions, however they are not necessarily conserved in their primary sequence nor in the structure of their precursors. The miR-200 family comprises five members that have been linked to T2D, causing β-cell mass loss by triggering apoptosis and decreasing insulin production (18). Thus, inhibitors of these miRs could serve as chemical probes to study their function and potentially as treatments of T2D if properly developed. Across the miR-200 family, miR-200c has been suggested as a driver of β-cell mass reduction (**Figure 1A**). Thus, we sought to design ligands to specifically inhibit this target by recognition of the three-dimensional structure of pre-miR-200c. By targeting this member of the miR-200 family specifically, we open the way for further investigation into the role of miR-200c in β-cell mass regulation.

We analyzed the sequences and structures of all miR-200 family members, which are shown in **Figure 1A & B** (19, 20). Not surprisingly, a high degree of sequence homology was observed between the mature 3p miRNAs, ranging from 72% to 95%, suggesting that sequence-based recognition would not be specific enough to inhibit a singular miR. To test if this is indeed the case, we obtained a phosphorothioate antagomir with locked nucleic acid (LNA) modifications complementary to 21 nucleotides of mature miR-200c-3p from Qiagen (**Figure 1B**) and studied its effect on the levels of all mature miR-200 family members by RT-qPCR in a validated cellular model of T2D, the mouse pancreatic β cell line MIN6 forced to express pre-miR-200c (18). [Note: the mouse and human miR-200 family are highly conserved both from a sequence and structural perspective (**Figure S1**). Sequence similarity ranges from 54 – 98%. The antagomir oligonucleotide is 67 – 100% complementary to the family members, but fully complementary to miR-200c-3p (**Figure S1**).] Not unexpectedly, the oligonucleotide could not discriminate between the family members in the mouse pancreatic β cell line MIN6 at any concentration tested (0.1 – 100 nM; **Figure 1C**), even with forced expression of miR-200c and endogenous levels of the others.

In contrast with the high degree of sequence homology between miR-200 family members, significant differences in the structures of their precursors were observed (**Figure 1B**; see **Figure S1** for a comparison of the structures of mouse and human miRs). This suggests that a structure-targeting ligand may be able discriminate amongst the family members. Our previous studies have shown that binding of small molecules to functional sites in miRs, *i.e.*, Drosha and Dicer processing sites (7, 21), can impede miRNA biogenesis, de-repress downstream proteins, and rescue disease phenotype; small molecules that bind to other sites in the miRNA are inactive (5, 22). Fortuitously, our analysis also revealed that the 1×1 nucleotide UU Internal loop in pre-miR-200c’s Dicer site is unique across the family and thus is an ideal target site against which to develop a specific ligand. [Note the structural similarity between mouse and human pre-miR-200c, including the Dicer site and adjacent regions (**Figure S1**).]

We queried our database of RNA structure-small molecule interactions to identify lead small molecules for the UU internal loop in miR-200c’s Dicer processing site, *i.e.*, our lead identification strategy Inforna (11, 23). Inforna identified a promising lead in compound **1** (**Figure 2**), which bound avidly to this structure with a K_d_ of 1.1±0.08 µM. The compound had no measurable affinity to an RNA with a single nucleotide change that alters the 1×1 nucleotide UU internal loop to a base pair (**Figure S2**). *In vitro*, **1** inhibited Dicer processing of pre-miR-200c with an IC_50_ of 2.2±0.5 µM, with no effect on processing of a control pre-miR-200c RNA in which the binding site for **1** was mutated to an AU base pair (**Figure S3**). In MIN6 cells, **1** modestly reduced mature miR-200c levels (∼35% knock down at a dose of 20 μM) but was selective across the miR-200 family (**Figure S4**). Further, the reduction in mature miR-200c levels by **1** was dose-dependent (**Figure S5A**); in agreement with its designed mode of action, **1** also boosted levels of pre-miR-200c, by ∼50% at a 20 μM dose (**Figure S5B**). As a result of the inhibition of pre-miR-200c biogenesis, **1** de-repressed the miR’s downstream mRNA targets *Rps6kb1* (ribosomal protein S6 kinase β-1 isoform), *Dnajc3* (DnaJ Heat Shock Protein Family (Hsp40) Member C3), *Xiap* (X-linked inhibitor of apoptosis), and *Jazf1* (juxtaposed with another Zinc finger protein 1) (**Figure S5C**), and inhibited β-cell apoptosis, a miR-200c-associated phenotype in T2D (**Figure S5D**).

**Figure 2:**
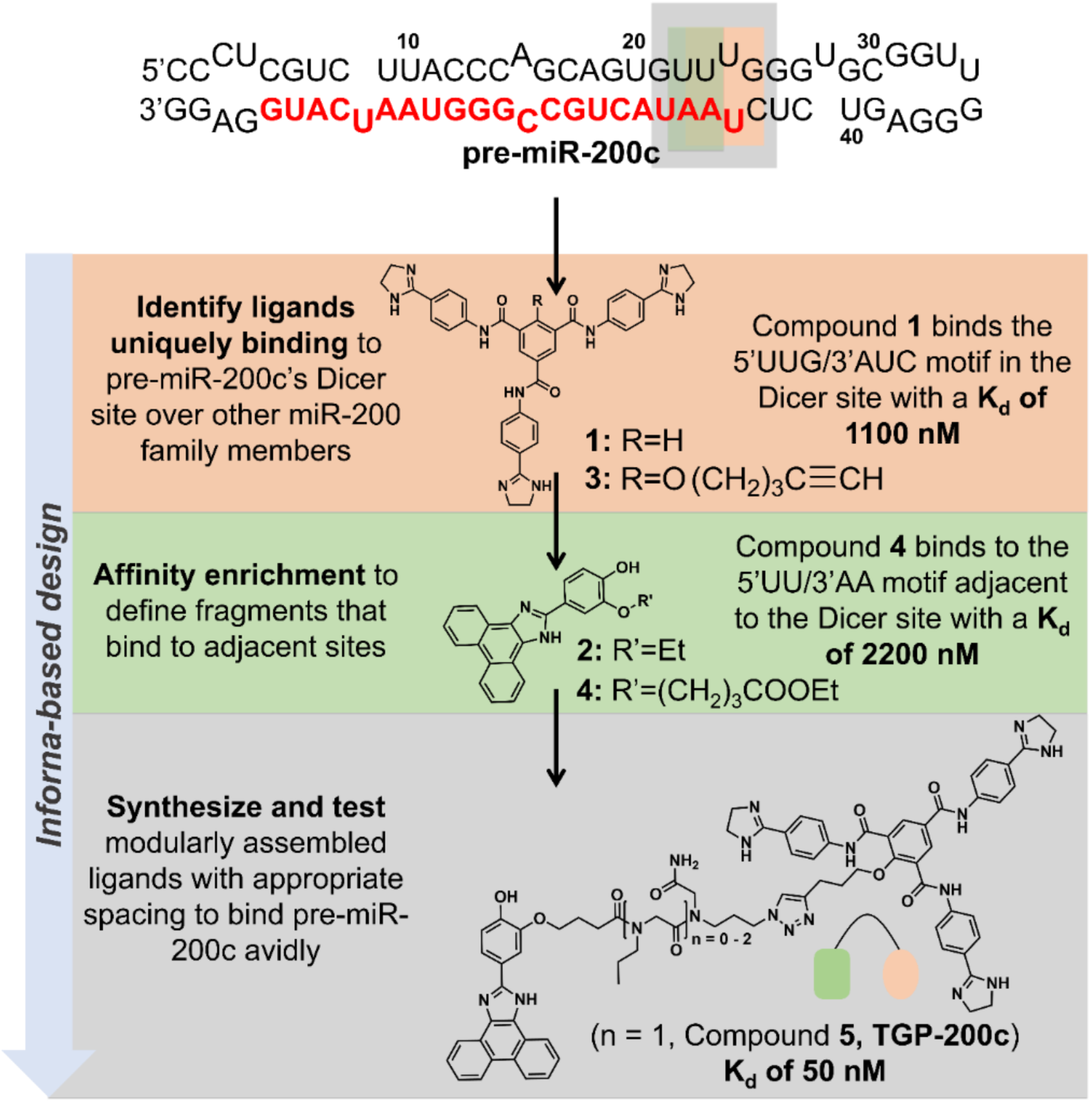
Inforna-based design of small molecules targeting pre-miR-200c. The secondary structure of pre-miR-200c was mined via Inforna, identifying compound **1** as a binder to the U/U loop at the Dicer processing site (K_d_ =1100 nM). Further analysis identified compound **2** that binds to the adjacent AU base pair (K_d_ = 2200 nM). Linking these two monomers (via **3** and **4**) afforded a more avid binder to the Dicer site of pre-miR-200c, **TGP-200c** (K_d_ = 50 nM).

### Inforna-based optimization of 1 increases in vitro affinity and potency

Given that **1** binds to the 1×1 nucleotide UU internal loop at the Dicer site in pre-miR-200c with modest affinity and inhibits pre-miR-200c with good selectivity in pancreatic β cells, it provides a promising lead for further optimization. One way to increase affinity, selectivity, and potency is to bind two or more structural motifs simultaneously with a single small molecule. Typically, we have designed modularly assembled compounds that target two adjacent non-canonically paired motifs, such as internal loops that are in close proximity (24). In pre-miR-200c, however, such sites are distant from the binding site of **1**. Thus, we designed a heterodimer comprising **1** and ligand **2** that binds a 5’UU/3’AA base pair motif adjacent to the Dicer processing site with high affinity and selectivity (**Figure 2**) (25).

Synthesis of the heterodimer required modification of **1** and **2** with functional handles for conjugation. Specifically, **1** was functionalized with an alkyne for azide-alkyne click chemistry (26), affording **3**, while **2** was functionalized with a butanoic acid linker for subsequent coupling to an amine, affording **4**, respectively. Analysis of **3** and **4** showed that they have similar *in vitro* and biological activity compared to their respective parent compounds **1** and **2** (**Figures S6 - S8**). [Note: **2** and **4** did not inhibit Dicer processing of pre-miR-200c *in vitro* or in MIN6 cells, as expected as it does not bind the Dicer processing site, but did bind pre-miR-200c with modest affinity.] The RNA-binding modules were assembled onto peptoid scaffolds of different lengths (n = 0, 1, and 2 spacing modules) to identify the one that most closely mimics the distance between pre-miR-200c’s Dicer site and adjacent AU pairs (**Figure 2**). The most promising dimer was identified by screening the three dimers for inhibiting apoptosis of MIN6 cells at 0.02, 0.2, 2 μM concentrations. While dimers with 0 and 2 spacing modules were inactive, the dimer with one *N*-propyl glycine spacing module, **5**, exhibited dose dependent rescue of apoptosis (reduction of Caspase 3/7 activity; **Figure S9A & B**). Notably, **TGP-200c** was non-toxic to MIN6 cells at these concentrations (**Figure S9C**). Heretofore, **5** is referred to as **TGP-200c** (Targapremir-200c) as it selectively and potently targets pre-miR-200c *vide infra*.

We next studied **TGP-200c** *in vitro* and compared it with lead compound **1. TGP-200c** bound pre-miR-200c with a K_d_ of 50±8 nM, 22-fold more avidly than **1** (**Figure S6**). Furthermore, no binding was observed when either the 1×1 nucleotide UU internal loop was mutated to an AU base pair or when the 5’UU/3’AA motif was mutated to GC pairs (**Figure S6**). Thus, both the internal loop and the adjacent 5’UU/3’AA are required for binding of **TGP-200c. TGP-200c**’s enhanced affinity translated into enhanced potency *in vitro*, inhibiting Dicer processing of pre-miR-200c with an IC_50_ of ∼200 nM nM, 10-fold more potent than **1** (**Figures S3A & S10A**). Importantly, **TGP-200c** had no inhibitory effect on Dicer processing of a mutant pre-miR-200c in which the binding site for **1** was ablated (**Figure S10B**). Collectively the optimized assembled compound has improved binding affinity and potency relative to starting monomer **1** and is highly selective.

### TGP-200c occupies target pre-miR-200c in cells to a greater extent than 1

Chemical Cross-Linking and Isolation by Pull-down (Chem-CLIP) is a target validation method that can also be used to study target occupancy and selectivity *in vitro* and in cells (27). To study occupancy of pre-miR-200c, a Chem-CLIP probe was designed and synthesized. The probe, **6**, was constructed by using the alkyne functional group in **3** to attach a chlorambucil (CA) cross-linking module and a biotin module that allows for purification of cross-linked material. In MIN6 cells transfected with WT pre-miR-200c, probe **6** cross-linked with pre-miR-200c dose dependently (**Figure 3B**). Importantly, **6** did not bind a pre-miR-200c mutant in which the UU loop binding site for **1** was mutated to AU base pair in MIN6 cells, as expected due to its lack of binding affinity (**Figures 3B & S6B**).

**Figure 3:**
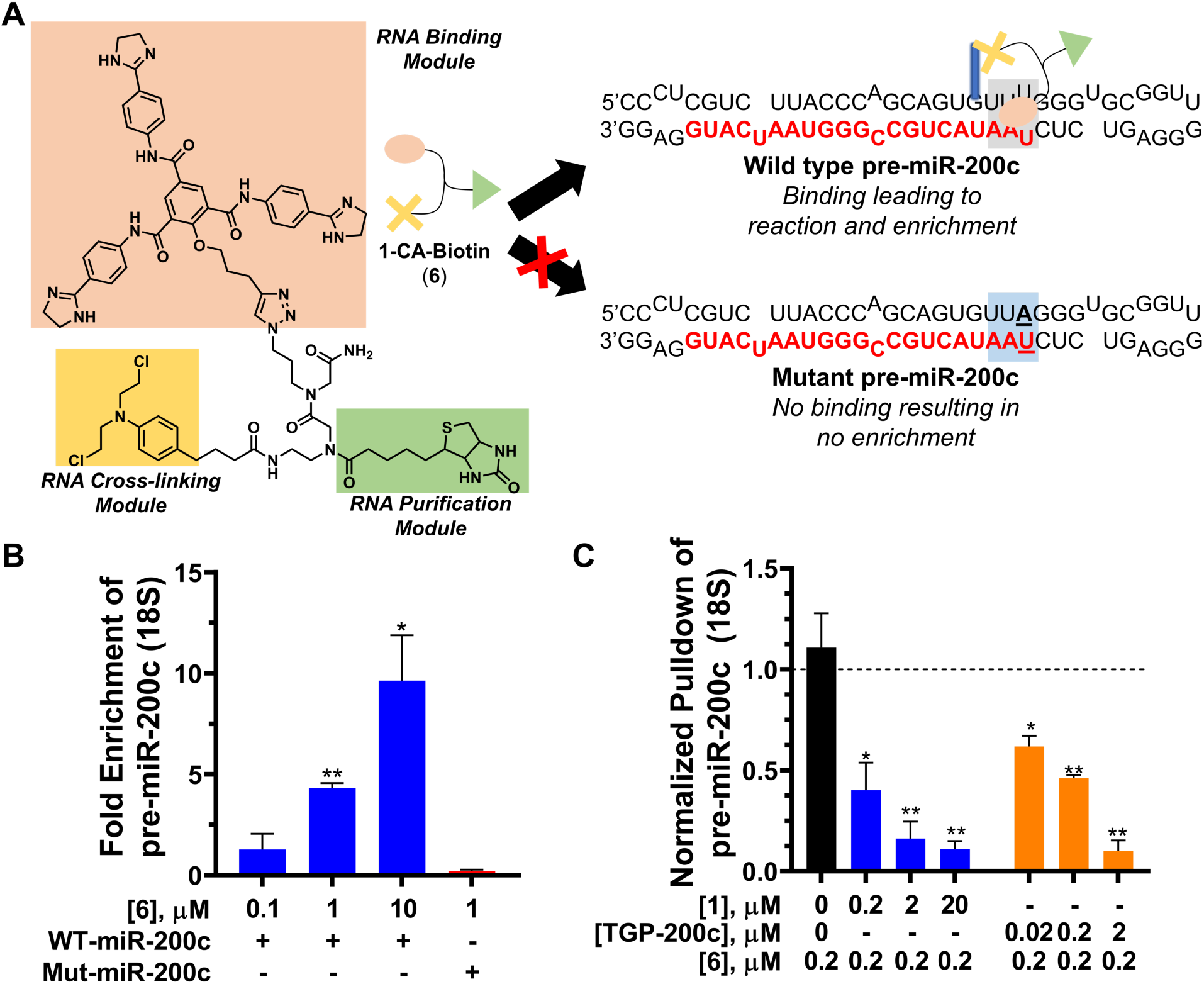
1 and TGP-200c selectively engage pre-miR-200c in cells. **A**) Scheme of Chemical-Cross Linking and Isolation by Pulldown (Chem-CLIP) showing that **6** binds and cross-links to wild type pre-miR-200c, while not engaging mutant pre-miR-200c. **B**) Chem-CLIP studies in MIN6 cells show dose dependent enrichment of pre-miR-200c by **6** for wild type miR-200c (WT-miR-200c), but not mutant miR-200c (Mut miR-200c) in which the binding is ablated (n = 3). **C**) Competitive-Chem-CLIP shows that **1** and **TGP-200c** can compete off the interaction of **6** with pre-miR-200c and, hence cross-linking. Note the 10-fold lower concentrations used for **TGP-200** than for **1** (n =3). *, p<0.05; **, p<0.01. All p-values were calculated by a two-tailed Student t-test. All data are reported as mean ± S.E.M.

To assess the relative occupancy of monomer **1** and **TGP-200c**, we used Competitive (C-)Chem-CLIP (5). In this experiment, a constant concentration of the Chem-CLIP probe **6** (200 nM) was added to cells along with increasing doses of unreactive **1** or **TGP-200c** (20 nM to 20,000 nM), which compete off the reaction of **6** with pre-miR-200c. Both compound **1** and **TGP-200c** inhibited cross-linking and hence pull down of pre-miR-200x by the Chem-CLIP probe. As expected, **TGP-200c** was a 10-fold more effective competitor than **1** (**Figure 3C**). All together, *in vitro* binding analyses, inhibition of Dicer processing *in vitro*, and cellular target occupancy studies suggest that the lead optimized compound more potently targets pre-miR-200c than starting compound **1**.

### TGP-200c potently inhibits miR-200c biogenesis selectively across the miRnome

We first measured **TGP-200c**’s ability to inhibit the miR’s biogenesis by measuring its effect on mature miR levels, which should decrease, and pre-miR levels, which should increase. A dose-dependent (0.02, 0.2, 2 μM) decrease in mature miR-200c levels was indeed observed with an IC_50_ of ∼200 nM, ∼300-fold more potently than **1** (**Figures 4A & S5A**). Importantly, even at the high dose of 2 μM, **TGP-2ooc** had no effect on the other members of the miR-200 family (**Figure 4B**), in contrast to the sequence-based recognition of the oligonucleotide antagomir (**Figure 1C**). Consistent with its designed mode of action, **TGP-200c** increased pre-miR-200c levels (**Figure 4C**). To assess the selectivity of **TGP-200c** for inhibiting miR-200c biogenesis, we studied its effects across all 193 miRs that are detectable in MIN6 cells by RT-qPCR. Indeed, only miR-200c levels were affected significantly, further demonstrating that the compound is potent and selective (**Figure 4D**).

**Figure 4:**
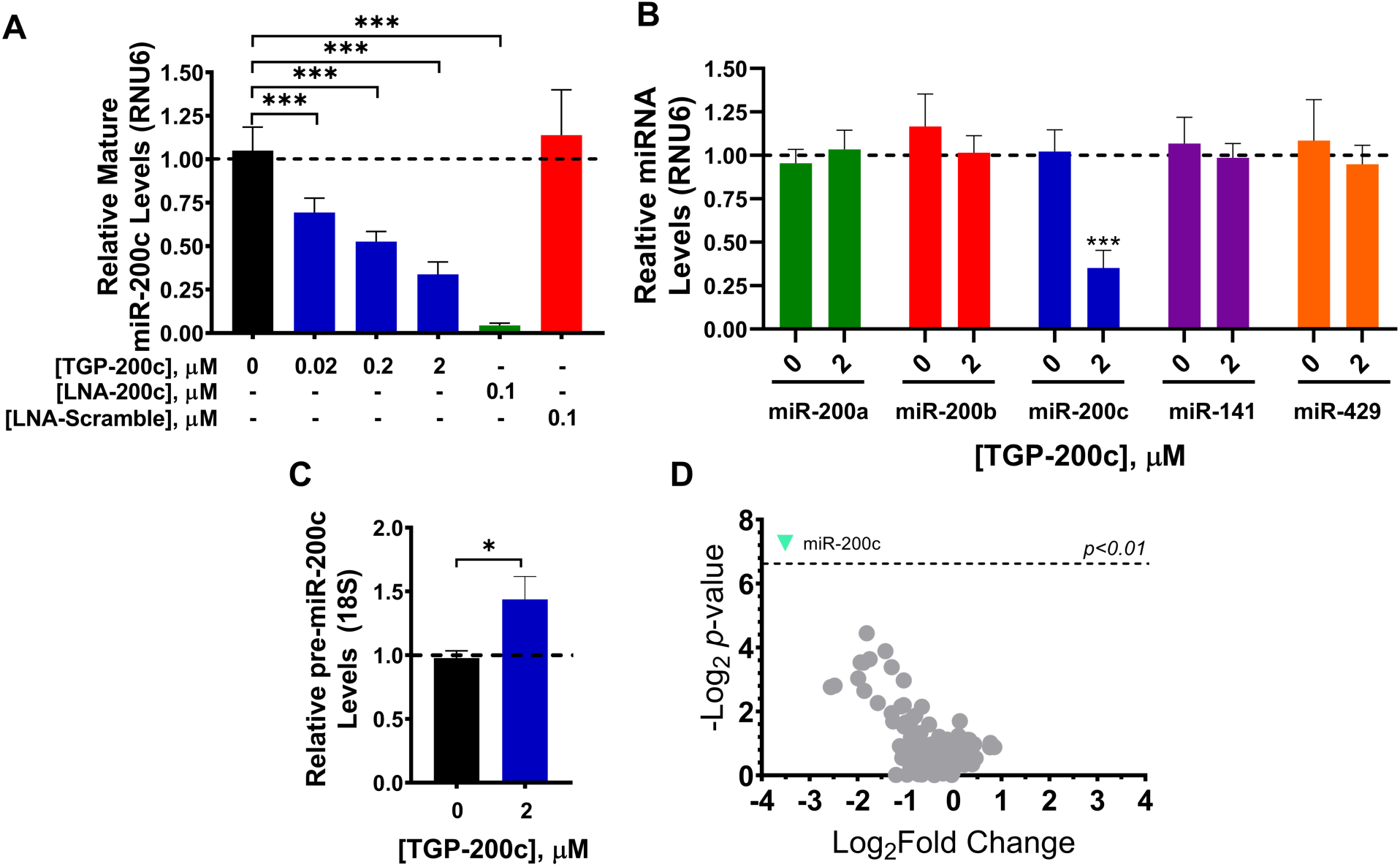
TGP-200c selectively inhibits miR-200c biogenesis in MIN6 cells. **A**) Effect of **TGP-200c** on mature miR-200c levels (n = 5). **B**) Effect of **TGP-200c** on mature levels of all miRs in the miR-200 family (n = 4). **TGP-200c** selectively inhibits miR-200c over all miR-200 family members by recognizing the U/U internal loop unique to pre-miR-200c. **C**) Effect of **TGP-200c** on pre-miR-200c levels (n = 4). **D**) Global profiling of all miRs expressed in MIN6 cells shows that only miR-200c levels are significantly downregulated by **TGP-200c** (p <0.01; n = 4). *, p<0.05; **, p<0.01. ***, p<0.001. All p-values were calculated by a two-tailed Student t-test. All data are reported as mean ± S.E.M.

As mentioned above, in T2D miR-200c acts by repressing the expression of *XIAP, RPS6KB1, DNAJC3*, and *JAZF1* by binding to their 3’ UTRs (**Figure 1**) (18). Thus, we measured **TGP-200c**’s ability to stimulate their expression levels by RT-qPCR. At a 2 μM dose, **TGP-200c** increased levels of each mRNA (**Figure 5A**). Robust antibodies are available for DNAJC3 and RPS6KB1, and thus we measured changes in their protein levels upon treatment with 2 μM of **TGP-200c** by Western blotting. Indeed, we observed an increase in the levels of both proteins by ∼40%, similar to the antagomir directed at miR-200c (**Figure 5B**).

**Figure 5:**
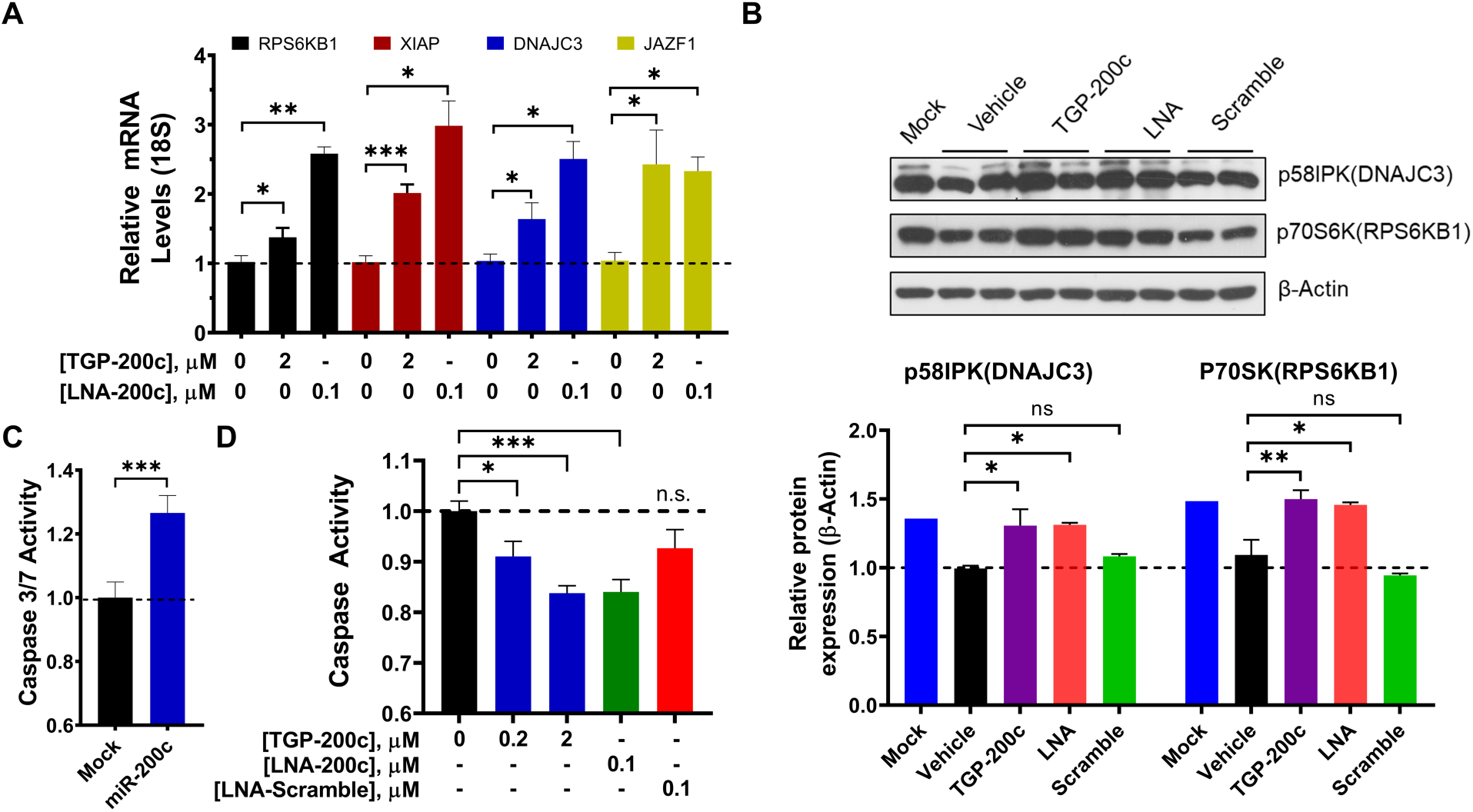
TGP-200c selectively rescues the miR-200c-mediated phenotype in MIN6 cells. **A**) Effect of **TGP-200c** on miR-200c’s direct mRNA, *Rps6kb1, Dnajc3, Xiap* and *Jazf1*, as determined by RT-qPCR (n = 4). **B**) Effect of **TGP-200c** on p58IPK (DNAJC3) and p70s6K (RPS6KB1) protein levels (n = 1 for Mock, n=4 for Vehicle and TGP-200c, n=2 for LNA and scramble). **C**) Effect of overexpression of pre-miR-200c on the apoptosis of MIN6 pancreatic β-cells, as determined by measuring Caspase 3/7 activity (n = 16). **D**) Effect of **TGP-200c** on Caspase 3/7 activity (n = 8). *, p<0.05; **, p<0.01; ***, p<0.001. All p-values were calculated by a two-tailed Student t-test. All data are reported as mean ± S.E.M.

In T2D, aberrant expression of miR-200c triggers apoptosis (**Figure 5C**), an effect that can be directly traced to decreased levels of Xiap and Rps6KB1 (**Figure 1**). De-repression of these proteins by **TGP-200** suggests that the compound may rescue this T2D phenotype. As assessed by Caspase 3/7 activity, **TGP-200c** inhibited apoptosis on MIN6 cells transfected with pre-miR-200c dose dependently (**Figure 5D**). Thus, **TGP-200c** deactivates a T2D circuit at the level of the transcriptome, proteome, and phenotype.

To confirm the binding site dependence on the bioactivity of **TGP-200c**, we studied its effect on MIN6 cells expressing the mutant pre-miR-200c (mut-miR-200c) in which the compound’s binding site is ablated; the mutant is processed by Dicer but is not inhibited by **TGP-200c** (**Figure S11**). As expected, **TGP-200c** was unable to inhibit the biogenesis of mut-miR-200c, de-repress *Xiap, Rps6kb1, Dnajc3* and *Jazf1* mRNA levels, or inhibit apoptosis, each in contrast to the LNA antagomir (**Figure S11**). These studies demonstrate that **TGP-200c** exerts its specific effects by binding the Dicer site in pre-miR-200c and inhibiting its biogenesis.

### TGP-200c exerts selective effects on proteome globally

We next studied the selectivity of **TGP-200** on the entire proteome, dosing MIN6 cells transfected with pre-miR-200c with 2 μM compound. Only 13 of the 4000 detectable proteins were affected, using a false discovery rate (FDR) of 1%. Of these 13 proteins, seven are encoded by mRNAs that are direct targets of miR-200c/200b/429 which form one of two subfamilies within the miR-200 family (**Figure 6A**). In contrast, no significant effects are found on proteins whose mRNAs are targets for the miR-200a/miR-141 subfamily (**Figures 6A & 6B**).

**Figure 6.**
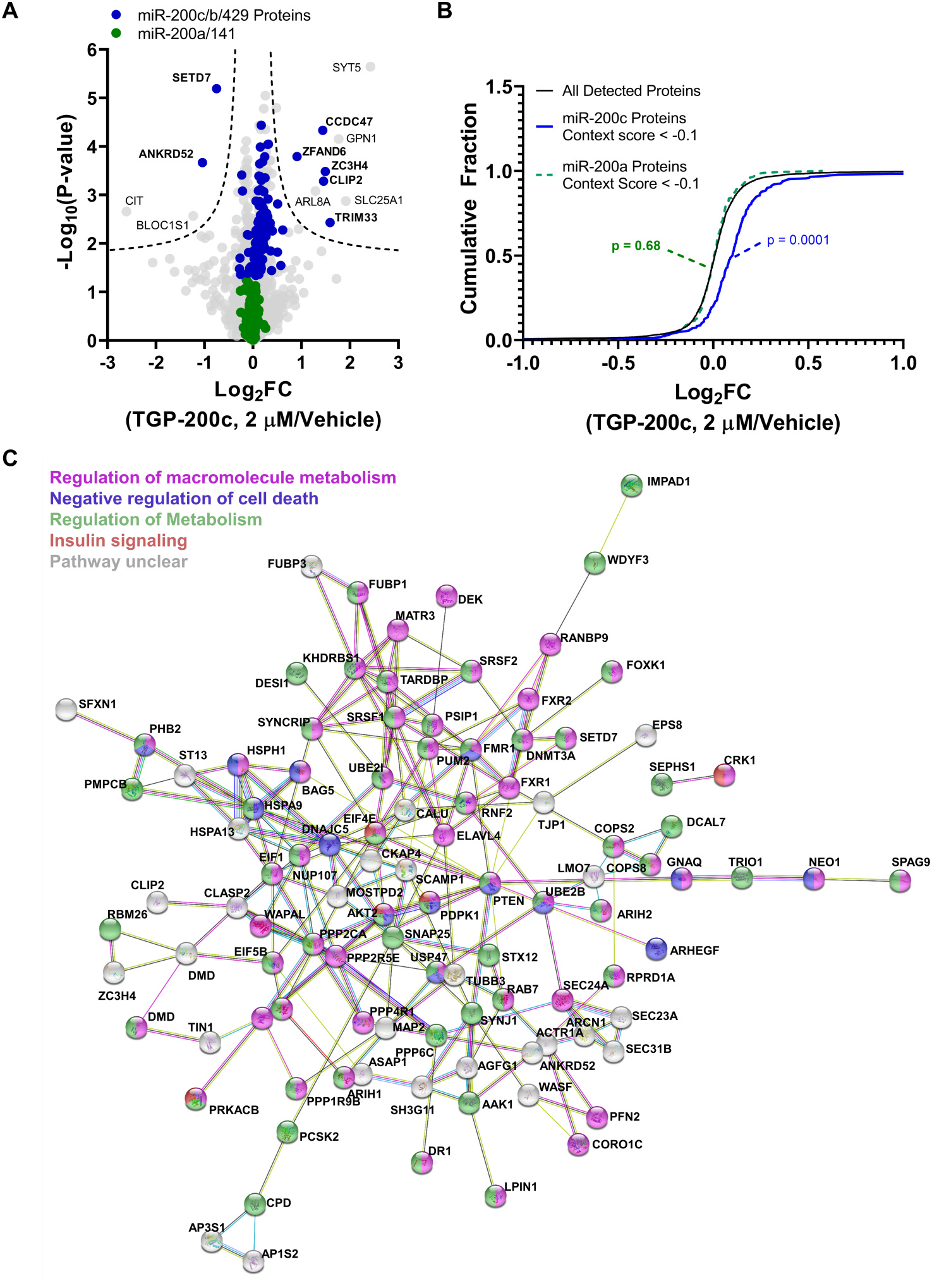
TGP-200c has specific effects on the global MIN6 proteome, traced to the selective inhibition of miR-200c biogenesis. **A**) Effect of **TGP-200c** (2 µM) on the proteome of MIN6 cells. Proteins encoded by mRNAs that are direct targets of miR-200c, −200b, and −429 (a miR-200 subfamily) are highlighted in blue; direct targets of miR-200a and miR-141 are highlighted in green. Downstream targets were predicted by TargetScan (48). **B**) On-target effects of **TGP-200c** observed in global proteomics analysis, grouped by miR-200 subfamilies, miR-200c/200b/429 and miR-200a/141 protein. The difference in their downstream targets arises from the single nucleotide difference in their seed regions (U/C in the third position). **C**) Pathway analysis of 128 proteins predicted to be regulated by miR-200c/-200b/-429 conducted using STRING analysis (www.string-db.org). Several pathways are selectively modulated by **TGP-200**, including insulin signaling (red), macromolecule metabolism (pink), general cellular metabolism (green), and negative regulation of cell death (purple) with false discovery rates of 0.025, 2.3×10^−8^, 0.004, and 0.0009, respectively.

Pathway analysis revealed that various T2D-related pathways are modulated by **TGP-200c** treatment, including insulin signaling and metabolism (**Figure 6C**). Interestingly, we see an upregulation of proteins that negatively regulate cell death such as AKT2 (28) among others, which is also expected given the anti-apoptotic phenotype of **TGP-200c**. These observations also support our hypothesis that **TGP-200c** represses pro-cell death factors and promotes β-cell survival in specific ways. Collectively, these proteome-wide studies and pathway analyses further bolster the assertion that **TGP-200c** is selective small molecule.

## Discussion

The most common way to target RNAs is by the sequence-based recognition of oligonucleotides (4). RNA 3D-structure, however, directly influence and in some cases control biology (29). Thus, targeting RNA structures with small molecules to perturb their function is a route to affect and study RNA biology (3). There is the perception that human RNAs do not generally fold into well-defined structures. Coupled with the notion that RNA cannot form diverse structures as it is only built on four nucleic acid building blocks, RNA has been mostly thought of as “undruggable” or “unligandable”. A variety of studies have challenged this assertion, including those presented herein. Evidently, RNA is indeed a viable small molecule drug target and targeting RNA structures in cells can be done rationally and predictably to provide potent and selective small molecules across a variety of indications (27), now including T2D.

Oligonucleotides are thought to be selective for fully complementary targets, however they can have off-targets (30). These off-targets include RNAs that are fully complementary and those that contain mismatches (30-32). Other factors that affect selectivity include the expression levels of the on- and the off-targets, with the more highly expressed target more likely to be occupied and affected (32).

Previously, we have shown that for some RNA targets, structure-specific recognition can be advantageous in disease-affected cells and animal models over sequence-specific recognition (33-36). For example, r(CUG) repeat expansions that cause myotonic dystrophy type 1 form a robustly folded structure that can be targeted selectively with structure-binding small molecules over transcripts with short, unstructured stretches of r(CUG) repeats (37). In contrast, oligonucleotides complementary to the repeats are unable to distinguish between repeats of length that cause disease and those that do not. In line with those studies, herein we have shown that miR family members sharing sequence homology cannot be selectively inhibited with an oligonucleotide but the structures of their precursors can be distinguished by small molecules, resulting in a selective inhibitor of a single miR’s biogenesis as demonstrated by miRnome- and proteome-wide studies. That is, the small molecule exploits a structure unique to pre-miR-200c amongst its family members, and it exerts specific effects on the transcriptome, proteome, and on phenotype. This selective probe molecule is further believed to be a useful tool to disentangle the biological importance of the individual members in the miR-200 family beyond β-cell apotosis. The miR-200 family is one of the most well studied families within cancer (38), and miR-200c was recently reported to have a relevance in podocyte dysfunction in kidney disease (39).

Interestingly, our designed inhibitor comprises an internal loop-targeting small molecules and a base pair-targeting small molecule. There are limited examples of using such a strategy to target RNA as most dimeric molecules target two non-canonically paired structures (40). There could be a perception that adding a base-pair targeting module would negatively impact selectivity, but the variety of studies shown here with **TGP-200c** demonstrate that this is not necessarily the case. There are likely many more applications of such an approach, which could provide an efficient route to ligands that target RNA in which the linker size and hence molecular weight of the molecule could be reduced.

More broadly, T2D affects a large segment of the world’s population and there are very few therapeutic approaches (41). The ability to provide a lead compound that targets an ncRNA potently and selectively to abrogate a pro-apoptotic pathway suggests that these compounds could have utility as leads for further preclinical development. The data presented herein further cement that RNA is indeed targetable with small molecule broadly and across disease indications.

## Methods

### General Methods

All RT-qPCR experiments were completed on an Applied Biosystems QS5 384-well qPCR system. Gene expression was measured using Power Sybr Green Master mix (Life Technologies) for all RT-qPCR experiments. Gels with radioactive signal were imaged with a Typhoon FLA9500 (GE Healthcare Life Sciences), and images were processed using BioRad’s QuantityOne. RNA oligonucleotides used for binding assays were obtained from Dharmacon and deprotected according the manufacturer’s protocol. All RNAs were desalted on PD-10 sephadex columns (GE Healthcare Life Sciences) per the manufacturer’s protocol and quantified by UV/Vis on a DU800 UV-Vis spectrophotometer at 90 °C.

### Tissue Culture

MIN6 mouse pancreatic β cells (C0018008) were obtained from Addex Bio technologies and checked for mycoplasma contamination every 6 months by PCR assay (Promokine). Cells were maintained in growth medium: optimized DMEM from Addex Bio (catalog number C0003-02) supplemented with 20% (v/v) FBS (Sigma), 1× Penicillin/Streptomycin (Corning), and 35 mM of β-mercaptoethanol (Sigma). Growth medium was passed through a 0.2 µm sterile filter before use. All cells are cultured at 37 °C in 5% CO_2_.

### PCR amplification and transcription of DNA templates

DNA templates for pre-miR-200c and the mutant pre-miR-200c were PCR amplified using a forward primer that also encodes for a T7 RNA polymerase promoter and the corresponding a reverse primer (see **Table S1**). PCR amplification was carried out in 300 µL of 1× PCR buffer (10 mM Tris-HCl, pH 9.0, 50 mM KCl, and 0.1% (v/v) Triton X-100), 4.25 mM MgCl_2_, 0.33 mM dNTPs, 2 µM of each primer, and 1.5 µL of Taq DNA polymerase. Amplification was carried out by completing 30 cycles at 95 °C for 30 s, 50 °C for 30 s, and 72 °C for 60 s. Amplification was confirmed by a 3% (w/v) agarose gel stained with ethidium bromide.

*In vitro* transcription of the DNA templates was completed as reported previously (42). Briefly, 300 µL of the PCR product was incubated in 1 mL of 1× Transcription buffer (40 mM Tris-HCl, pH 8.0, 1 mM spermidine, 10 mM DTT, and 0.001% (v/v) Triton X-100) containing 2.5 mM of each rNTP, 15 mM MgCl_2_, and 20 µL of 20 mg/mL T7 RNA polymerase overnight at 37 °C. After transcription, 1 unit of RNase-free DNase I (Promega) was added to the reaction, which was incubated at 37 °C for an additional 1 h. The RNA was purified on a denaturing 15% polyacrylamide gel and extracted as previously described (42). RNAs were quantified by using its absorbance at 260 nm at 90 °C and the respective extinction coefficient determined from nearest neighbor parameters (43) and the HyTher server (44).

### Inhibition of Dicer processing of pre-miR-200c *in vitro*

Pre-miRs were radioactively labeled on the 5’ end and purified as previously described (42). The labeled RNA was folded in 1× Dicer Reaction Buffer (Genlantis) by heating at 95 °C for 30 s and cooling slowly to room temperature. The samples were then supplemented with 1 mM ATP and 2.5 mM MgCl_2_, followed by addition of **1** or **TGP-200c** (25 to 0.78 µM). The samples were incubated at room temp for 30 min followed by addition of recombinant Dicer (BPS-Bioscience) to a final concentration of 0.017 mg/mL. The samples were incubated for 1 h at 37 °C, and then the reaction was quenched by addition of an equal volume of 2× Loading Buffer (89 mM Tris-Borate, pH 8.3, 200 mM EDTA, 7M urea, 1 mg/mL bromophenol blue/xylene cyanole).

A T1 ladder (cleaves G residues) was generated by incubating 1 pmol of radioactively labeled RNA in 1× RNA Sequencing Buffer (20 mM sodium citrate, pH 5.0, 1 mM EDTA, and 7 M urea) at 95 °C for 30 s followed by slow cooling to room temperature. RNase T1 was then added to a final concentration of 3 U/µL, and the samples were incubated at room temp for 30 min. An RNA hydrolysis ladder was created by incubating 1 pmol of radioactively labeled RNA in 1× RNA Hydrolysis Buffer (50 mM NaHCO_3_, pH 9.4, and 1 mM EDTA) at 95 °C for 3 min followed by snap cooling on ice. An equal volume of 2× Loading Buffer was added to each sample.

The cleavage products were resolved on a 0.7 mm thick, denaturing 15% polyacrylamide gel and imaged using a Typhoon FLA 9500 by autoradiography.

### Microscale thermophoresis binding

The binding affinities of **1, 3, 4**, and **TGP-200c** were measured by microscale thermophoresis (MST). Measurements were taken using standard capillaries (Nano Temper) on a Monolith Nt. 115 system (Nano Temper). Samples were prepared with as follows: 20 nM of RNA was folded in 2× Binding Buffer (16 mM Na_2_PO_4_, pH 7.0, and 370 mM NaCl) by heating at 65 °C for 20 min followed by slow cooling to room temperature on the benchtop. Once cooled, bovine serum albumin (BSA) was added to a concentration of 80 µg/mL. Compounds were diluted to 100 µM and then serial diluted 1:2 in water. An equal volume of folded RNA has added to each concentration of compound in a total volume of 20 µL. The samples were incubated at room temp for 15 min before loading into the capillaries. Instrument settings were as follows: LED power 15%, MST power 40%, fluorescence detection before 5 s, MST power on 30 s, fluorescence detection after MST off 5 s, delay before next scan 25 s. Each run was repeated in quadruplicate, and the results averaged. All data were fit according to equation 1 below used previously(45): 

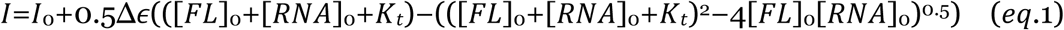

where I and I_0_ are the observed fluorescence and initial fluorescence intensity in the presence and absence of RNA, Δε is the difference between the fluorescence intensity in the absence and presence of infinite RNA concentration, [FL]_0_ and [RNA]_0_ are respectively the concentrations of the small molecule and RNA, and K_t_ is the dissociation constant.

### Transfection of MIN6 cells

MIN6 cells were grown to ∼50% confluency in 100 mm dishes and then batch transfected with 1 µg of wild type pre-miR-200c (MmiR-3304-MR04, Genecopoeia) or mutant pre-miR-200c (Synthesized by Genscript) using Lipofectamine 3000 (Life Technologies) per the manufacturer’s protocol. Cells were also mock transfected by using the same quantities of reagents for plasmid transfection without the addition of plasmid.

### Measuring pre- and mature miR-200c and mRNA levels by RT-qPCR analysis

After transfection, cells were allowed to recover for 8 h in complete growth medium before splitting into 12-well plates (100,000 cells/well). After adhering for 12 h, the cells were treated with **1** or **TGP-200c** in dose response, vehicle (DMSO), LNA-200c, or LNA-Scramble for 60 h. Total RNA was extracted using the Zymo Quick-RNA Miniprep Kit per the manufacturer’s protocol, including DNase treatment. Reverse ttranscription (RT) for miRs was completed with 300 ng of total RNA using a miScript II RT Kit (Qiagen) per the manufacturer’s protocol. RT of pre-miR-200c and mRNAs was completed using QScript Kit (Quanta Bio) with 1000 ng of total RNA per the manufacturer’s protocol. RT-qPCR was completed using a QS5 384-well PCR system (Applied Biosystems) using Power Sybr Green Master Mix and the manufacturer’s Comparative C_t_ with melt standard method, adjusted for a sample volume of 10 µL. All data were analyzed using the ΔΔC_t_ method previously described (46).

### Immunoblotting of RPS6KB1 and DNAJC3

Transfected MIN6 cells were seeded into 6-well plates at 200,000 cells/well and allowed to adhere for 12 h. After adhering, cells were treated with **1, TGP-200c**, LNA-200c, LNA-Scramble, or vehicle for 96 h. After 60 h, the medium was replaced with fresh medium containing compound. After 96 h, the cells were washed with 1× DPBS and then trypsinized and pelleted. The pellets were washed once with 1× DPBS before lysis in M-PER buffer (Thermo Fisher) supplemented with 1× protease inhibitor cocktail (Roche). Samples were subjected to a freeze-thaw cycle (−80 to room temperature) to ensure complete lysis. Total protein samples were quantified using a Pierce Micro BCA Protein Assay Kit (Fisher Scientific) per the manufacturer’s protocol. Approximately 20 µg of total protein weas resolved on a 10% SDS-polyacrylamide gel and then transferred to a PVDF membrane. The membrane was blocked in 5% (w/v) milk in 1× TBST (1× TBS, pH 7 supplemented with 0.1% (v/v) Tween 20) for 30 min at room temperature. The membrane was then incubated with 1× TBST containing 5% milk and either anti-RPS6KB1 (Cell Signaling Technology, 2940S) or anti-DNAJC3 (Cell Signaling Technology, 2708s) primary antibody (1: 2000) at 4 °C overnight. The membrane was washed 1× TBST three times for 10 min each followed by incubation with anti-rabbit IgG horseradish-peroxidase secondary antibody conjugate (1:3000 dilution; Cell Signaling Technology, 7074S) in 1× TBS for 2 h. After washing with 1× TBST (3 × 10 min), protein expression was detected using SuperSignal West Pico Chemiluminescent Substrate (Thermo Fisher) according to the manufacturer’s protocol.

The membrane was then stripped by incubating with 1× Stripping Buffer (200 mM glycine and 0.1% SDS, pH 2.2) for 1 h at room temperature. The membrane was washed with 1× TBS for 20 min and then incubated with anti-β-actin primary antibody (1:10000 dilution; Cell Signaling Technology, 3700S) in 1× TBST with 5% milk for 2 h at room temperature. The membrane was then washed with 1× TBST (3 × 10 min) and incubated with anti-mouse IgG horseradish-peroxidase secondary antibody conjugate (1:10000 dilution; Cell Signaling Technology, 7076S) in 1× TBST for 1 h at room temperature. After washing with 1× TBST (3 × 10 min), β-actin expression was detected as described above. ImageJ software was used to quantify the intensities of the protein bands in all Western blots.

### Cell viability and Caspase 3/7 Assay

Transfected MIN6 cells (15,000 cells per well) were seeded into 96-well white, clear bottom tissue culture plates (VWR: 89091-014) in 90 µL of growth medium and allowed to adhere for 12 h. The cells were then dosed with the compound or oligonucleotide of interest or vehicle and incubated for 60 h. Cell viability was measured using CellTiter Fluor (Promega) according to the manufacturer’s protocol, and fluorescence was measured using a Molecular Devices M5 plate reader.

To measure Caspase 3/7 activity, after the treatment period, 100 µL/well of Caspase 3/7 reagent (Promega) was added per the manufacturer’s protocol. Luminescence was measured on a Molecular Devices M5 plate reader, set to an integration time of 500 ms. Data were processed by first normalizing cell viability to vehicle controls and then normalizing the resulting data to Caspase 3/7 luminescence.

### Cellular Chem-CLIP and Competitive-Chem-CLIP of pre-miR-200c

MIN6 cells transiently expressing pre-miR-200c were treated with **1-CA-Biotin** (10, 1, and 0.1 µM) for 48 h in a 60 mm dish. After 48 h, total RNA was extracted using the miRNeasy Mini Kit (Qiagen) per the manufacturer’s protocol, with DNase I treatment. Then, 15 µg of total RNA in 500 uL of 1× DNA buffer (8 mM Na_2_PO_4_, pH 7.0, 185 mM NaCl) was incubated with 200 µL of a slurry of Dynabeads MyOne Streptavidin C1 (Invitrogen). The samples were incubated with shaking for 5 h at room temperature. The beads were then captured on a magnetic rack and washed with 1× Wash Buffer (10 mM Tris-HCl, pH 7.2, 1 mM EDTA, and 4 M NaCl) six times and once with nanopure water. The bound RNA was then released by incubating the beads with 150 µL of 95% formamide containing 10 mM EDTA at 95 °C for 30 min. The released RNA was then cleaned up by adding 2 volumes of RNA Lysis Buffer (Zymo Research) and one volume of 100% ethanol. The sample was loaded onto Quick RNA Miniprep columns (Zymo Research) and then proceeding according to the manufacturer’s protocol. RT-qPCR was carried out on pre-miR-200c as described above using 5 ng of input cDNA from before and after pull-down. Enrichment was calculated as previously described (22). C-Chem-CLIP was completed by pre-treating MIN6 cells with **1** (0.2, 2, 20 µM) or **TGP-200c** (0.02, 0.2, 2 µM) for 15 min followed by addition of 0.2 µM **1-CA-Biotin** for 48 h. Sample processing and analysis were carried out as described above.

### Global proteomics profiling using LC-MS/MS

Cells were harvested and suspended in 1× PBS and then lysed via sonication. Protein concentration was determined using a Bradford assay (BioRad). Samples (20 µg) were denatured with 6 M urea in 50 mM NH_4_HCO_3_ pH 8, reduced with 10 mM tris(2-carboxyethyl)phosphine hydrochloride (TCEP) for 30 min, and finally alkylated with 25 mM iodoacetamide for 30 min in the dark. Samples were diluted to 2 M urea containing 50 mM NH_4_HCO_3_ pH 8 and digested with trypsin (1 µL of 0.5 µg/µL) in the presence of 1 mM CaCl_2_ for 12 h at 37 °C. Samples were acidified with acetic acid to a final concentration of 5%, desalted over a self-packed C18 spin column, and dried. Samples were analyzed by LC-MS/MS (below) and the MS data were processed with MaxQuant (below).

### LC-MS/MS analysis

Peptides were resuspended in water with 0.1% formic acid (FA) and analyzed using EASY-nLC 1200 nano-UHPLC coupled to Q Exactive HF-X Quadrupole-Orbitrap mass spectrometer (Thermo Scientific). The chromatography column consisted of a 50 cm long, 75 µm i.d. microcapillary capped by a 5 µm tip and packed with ReproSil-Pur 120 C18-AQ 2.4 µm beads (Dr. Maisch GmbH). LC solvents were 0.1% FA in H_2_O (Buffer A) and 0.1% FA in 90% MeCN: 10% H_2_O (Buffer B). Peptides were eluted into the mass spectrometer at a flow rate of 300 nL/min over a 240 min linear gradient (5-35% Buffer B) at 65 °C. Data were acquired in data-dependent mode (top-20, NCE 28, R = 7’500) after full MS scan (R = 60’000, m/z 400-1’300). Dynamic exclusion was set to 10 s, peptide match to prefer, and isotope exclusion was enabled.

### MaxQuant analysis

MS data were analyzed with MaxQuant(47) (V1.6.1.0) and searched against the mouse proteome (Uniprot) and a common list of contaminants (included in MaxQuant). The first peptide search tolerance was set at 20 ppm; 10 ppm was used for the main peptide search; and fragment mass tolerance was set to 0.02 Da. The false discovery rate (FDR) for peptides, proteins and sites identification was set to 1%. The minimum peptide length was set to 6 amino acids, and peptide re-quantification, label-free quantification (MaxLFQ), and “match between runs were enabled. The minimal number of peptides per protein was set to two. Methionine oxidation was searched as a variable modification and carbamidomethylation of cysteines was searched as a fixed modification.

## Associated Content

### Supporting Information

The Supporting Information is available free of charge at DOI: [to be filled in]

The following data can be found in the Supporting information, (i) table of oligonucleotides used in this study; (ii) Supporting Information Figures 1-10; and (iii) synthetic methods and characterization of all compounds.

## Author Information

### Author Contributions

M.D.D. designed and directed the study; H.S.H. and X.L. designed and executed experiments; H.S.H conducted all cellular and binding experiments; X.L. synthesized and characterized all compounds used; D.A. and A.A. conducted global proteomics analysis of cellular samples; L.K. and M.L. provided critical intellectual feedback on experimental design.

### Data Availability

All raw data associated that support the findings of this study are available from the corresponding author upon reasonable request.

## Acknowledgments

This work was funded by the National Institutes of Health (R01 GM097455 to MDD). We would also like to thank Dr. Jessica L. Childs-Disney, Mr. Christopher Williams, and Dr. Bader Zarrouki (AstraZeneca) for their input and editing during the writing of the manuscript.

