## Supporting Appendix for "Small molecules exploiting structural differences within microRNA-200 precursors family members reverse a type 2 diabetes phenotype"

Figure S10: **TGP-200c** inhibits Dicer processing of wild type but not mutant pre-miR-200c ...14

Figure S11: **TGP-200c** does not inhibit mutant miR-200c and its associated downstream targets in MIN6 cells ...............................................................................................................15-16

Synthetic Methods ......................................................................................................... 17-38

References...................................................................................................................... 39

| **Table S1**: Sequences of oligonucleotides used in these studies. | | | |
|---|---|---|---|
| **Oligonucleotide** | **Sequence 5’ -> 3’** | **Experiment** | **Supplier** |
| miR-200a-3p | TAACACTGTCTGGTAACGATGT | RT-qPCR | IDT |
| miR-200b-3p | TAATACTGCCTGGTAATGATGA | RT-qPCR | IDT |
| miR-200c-3p | TAATACTGCCGGGTAATGATGGA | RT-qPCR | IDT |
| miR-141-3p | TAACACTGTCTGGTAAAGATGG | RT-qPCR | IDT |
| miR-429-3p | TAATACTGTCTGGTAAAACCGT | RT-qPCR | IDT |
| Pre-miR-200c Forward | TCCATCATTACCCGGCAGTA | RT-qPCR | IDT |
| Pre-miR-200c Reverse | CGTCTTACCCAGCAGTGT | RT-qPCR | IDT |
| DNAJC3 Forward | GGTGGACCTGCAGTACGAA | RT-qPCR | IDT |
| DNAJC3 Reverse | TGTCCGGCTGCGAGTAAT | RT-qPCR | IDT |
| RPS6KB1 Forward | GGTAGTCCACGAACACCTGTC | RT-qPCR | IDT |
| RPS6KB1 Reverse | GGTATTCCACAGGGGTCTGA | RT-qPCR | IDT |
| XIAP Forward | AAAAACACCATCGCTAACTAAAAA | RT-qPCR | IDT |
| XIAP Reverse | CTTGAAGCTAAATCCCATTCG | RT-qPCR | IDT |
| JAZf1 Forward | CCTCAGCTCCATGTGCAT | RT-qPCR | IDT |
| JAZF1 Reverse | GGTGACCATTCTTAGCATG | RT-qPCR | IDT |
| RNU6 Forward | ACACGCAAATTCGTGAAGCGTTC | RT-qPCR | IDT |
| Universal Reverse | GAATCGAGCACCAGTTACGC | RT-qPCR | IDT |
| 18S Forward | GTAACCCGTTGAACCCCATT | RT-qPCR | IDT |
| 18S Reverse | CCATCCAATCGGTAGTAGCG | RT-qPCR | IDT |
| Power LNA-200c | C*C*A*T*C*A*T*T*A*C*C*C*G*G*C*A* G*T*A*T*T* | RT-qPCR | Qiagen |
| Power LNA-Scramble | T*A*A*C*A*C*G*T*C*T*A*T*A*C*G*C* C*C*A* | RT-qPCR | Qiagen |
| Pre-miR-200c Forward | GGCCGGATCCTAATACGACTCACTATA GGCCCTCGTCTTACCCAGCA | Dicer Assay | IDT |
| Pre-miR-200c Reverse | CCTCCATCATTACCCGGC | Dicer Assay | IDT |
| Pre-miR-200c Template | CCCTCGTCTTACCCAGCAGTGTTTGGG TGCGGTTGGGAGTCTCTAATACTGCCG GGTAATGATGGAGG | Dicer Assay | IDT |
| Mutant pre-miR-200c Template | CCCTCGTCTTACCCAGCAGTGTT**A**GGG TGCGGTTGGGAGTCTCTAATACTGCCG GGTAATGATGGAGG | Dicer Assay | IDT |
| WT 200c Cy5 | Cy5-GCGUUUGGGGAAACUCUAAUGC | Binding Assay | Dharmacon |
| GC Pair Mutant Only 200c Cy5 | Cy5-GCG**GGU**GGGGAAACUC**UCC**UGC | Binding Assay | Dharmacon |
| U/U Mutant 200c Cy5 | Cy5-GCGUU**A**GGGGAAACUCUAAUGC | Binding Assay | Dharmacon |
| WT 200c Biotin | Bi-GCGUUUGGGGAAACUCUAAUGC | Binding Assay | Dharmacon |

| | | | |
|---|---|---|---|
| Double Mutant 200c Biotin | Bi-GCG**CCA**GGGGAAACUC**UGG**UGC | Binding Assay | Dharmacon |
| GC Pair Mutant Only 200c Biotin | Bi-GCG**GGU**GGGGAAACUC**UCC**UGC | Binding Assay | Dharmacon |
| U/U Mutant 200c Biotin | Bi-GCGUU**A**GGGGAAACUCUAAUGC | Binding Assay | Dharmacon |

* indicate phosphorothioate backbone modification; LNA nucleotides were not disclosed by the manufacturer
**Bold letters** indicate mutations in sequence.

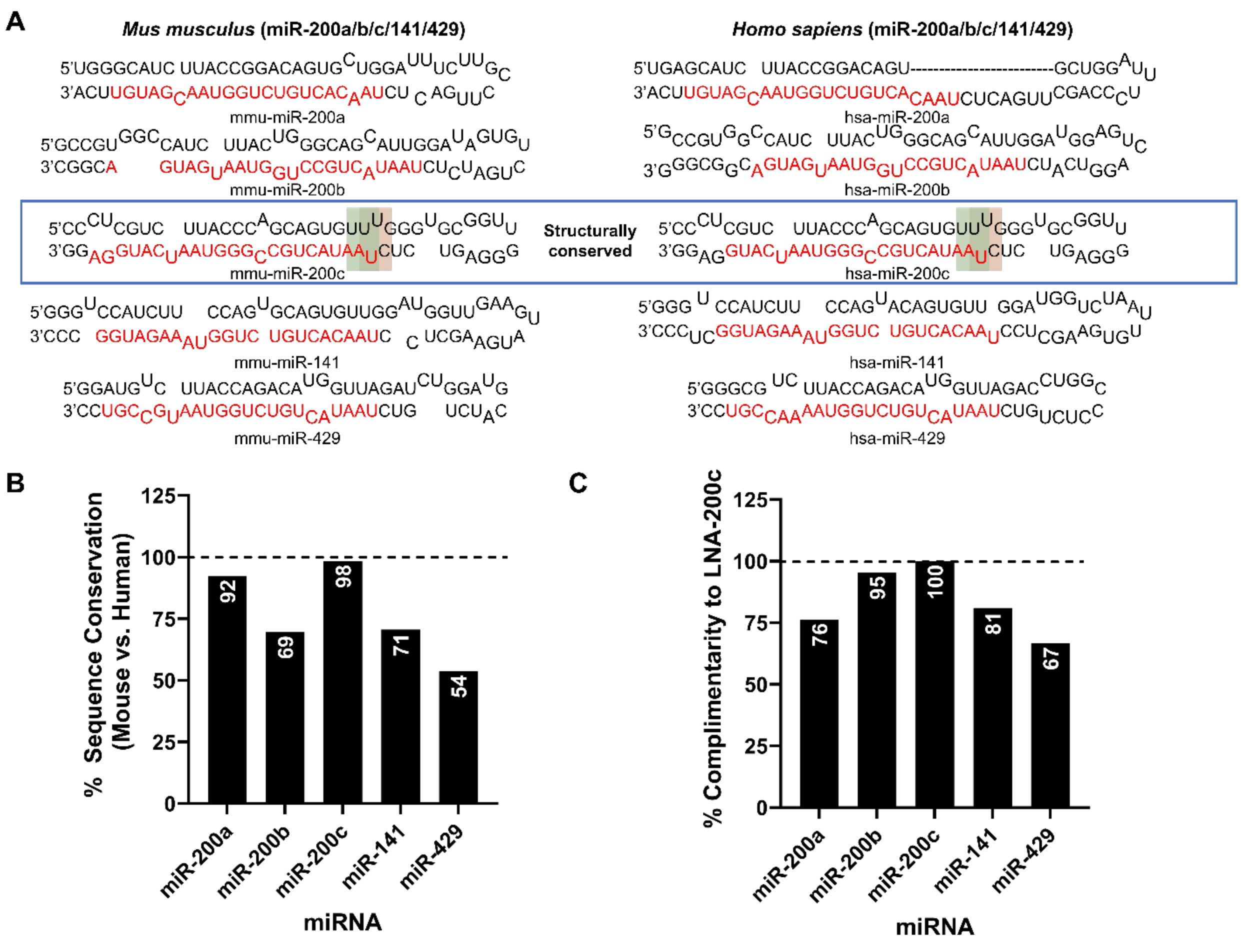

**Figure S1: Comparison of the sequences and structures of the miR-200 family from mouse and human.** **A**) Sequences of the mouse and human miR-200 family members. **B**) These comparisons show that miR-200c exhibits 98% sequence homology as well as structural homology between the mouse and human homologues whereas other family members do not. **C**) Complementarity of an LNA antisense oligo nucleotide targeting miR-200c shows diminished complementarity to other family members.

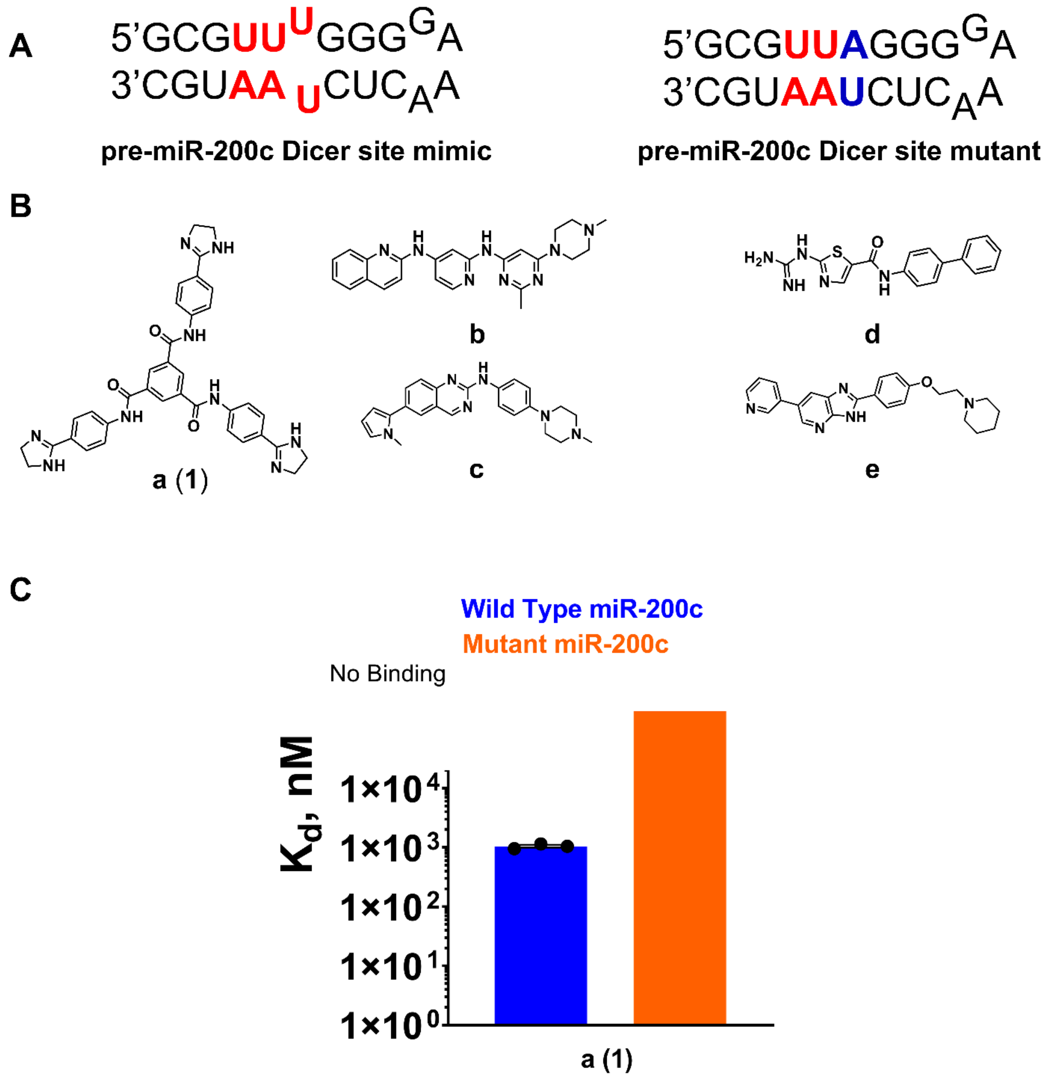

**Figure S2: Chemical structure of small molecules designed by Inforna to bind the Dicer processing site of pre-miR-200c and their binding affinities.** **A**) The secondary structures of RNAs used to measure binding affinity by microscale thermophoresis (MST). **B**) Chemical structures of the Inforna lead compounds for the UU internal loop present in pre-miR-200c's Dicer sitte. **C**) Binding summary of **1** binding to the miR-200c Dicer site (blue) and base paired control (orange) to RNA mimics. Compound a (**1**) exhibits selective binding to the Dicer site.

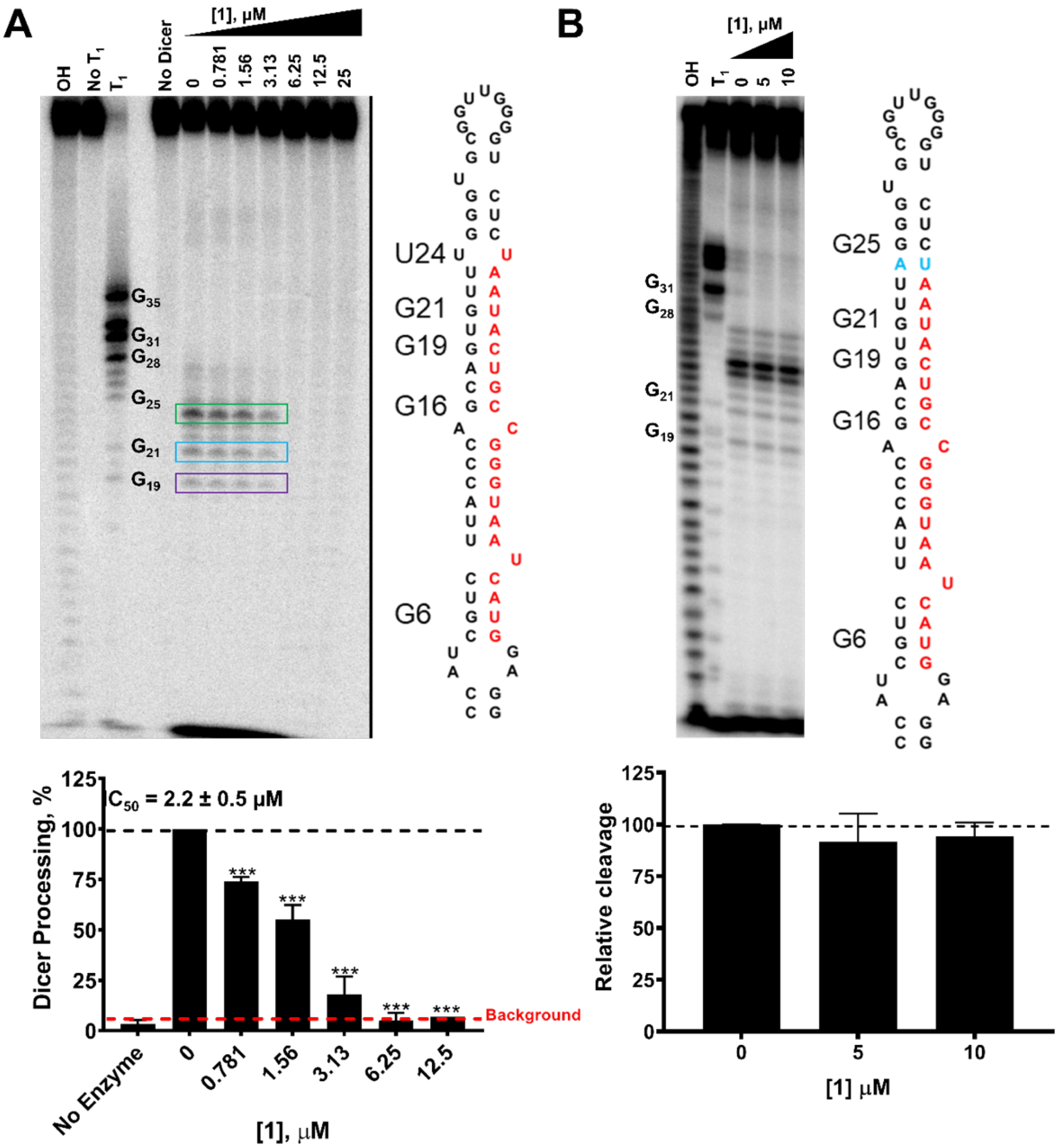

**Figure S3: *In vitro* Dicer inhibition of compound 1 to wild type and mutant miR-200c show that the compound only inhibits wild type.** **A**) Representative gel image of *in vitro* Dicer inhibition by **1**. Compound **1** inhibits Dicer processing with an $IC_{50}$ of 2.2 ± 0.5 μM, with as little as 780 nM showing 25% inhibition. **B**) *In vitro* Dicer processing of mutant pre-miR-200c shows that **1** has no effect on Dicer processing *in vitro* (*right*) (n = 3). ***, $p < 0.001$. All p-values were calculated by a two-tailed Student t-test. All data are reported as the mean ± standard error of the mean (S.E.M).

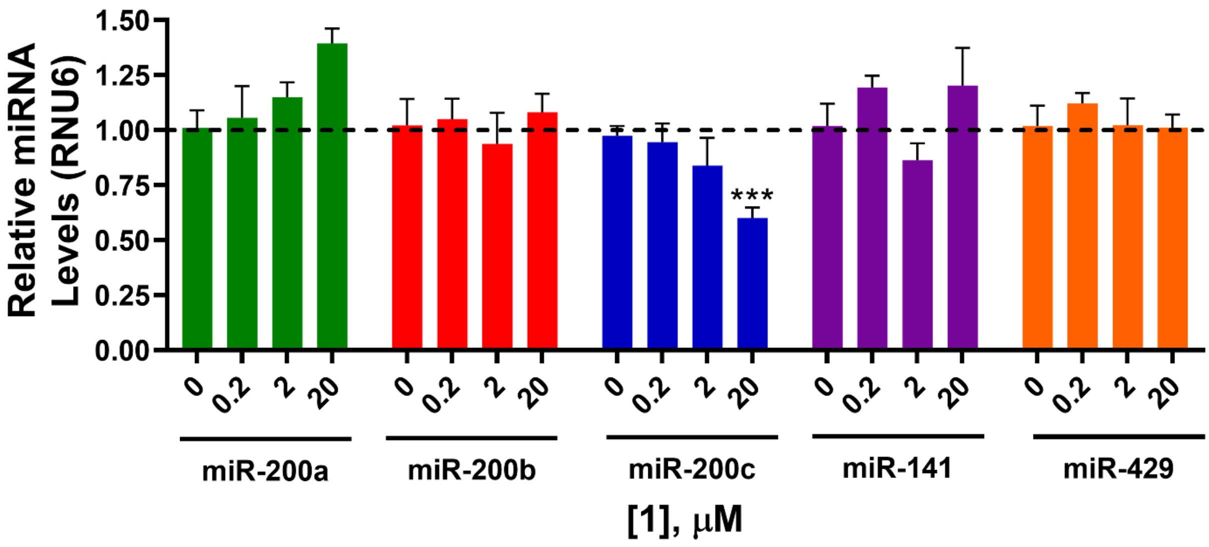

**Figure S4: Compound 1 selectively reduces mature miR-200c levels, as compared to all other miR-200 family members.** Compound **1** selectively reduced levels of mature miR-200c in MIN6 cells transfected with pre-miR-200c [induces downstream effects and an apoptotic phenotype consistent with T2D (1)]. Further, **1**'s reduction of miR-200c levels is dose dependent. ***, $p < 0.001$, as calculated by a two-tailed Student t-test. All data are reported as the mean ± S.E.M.

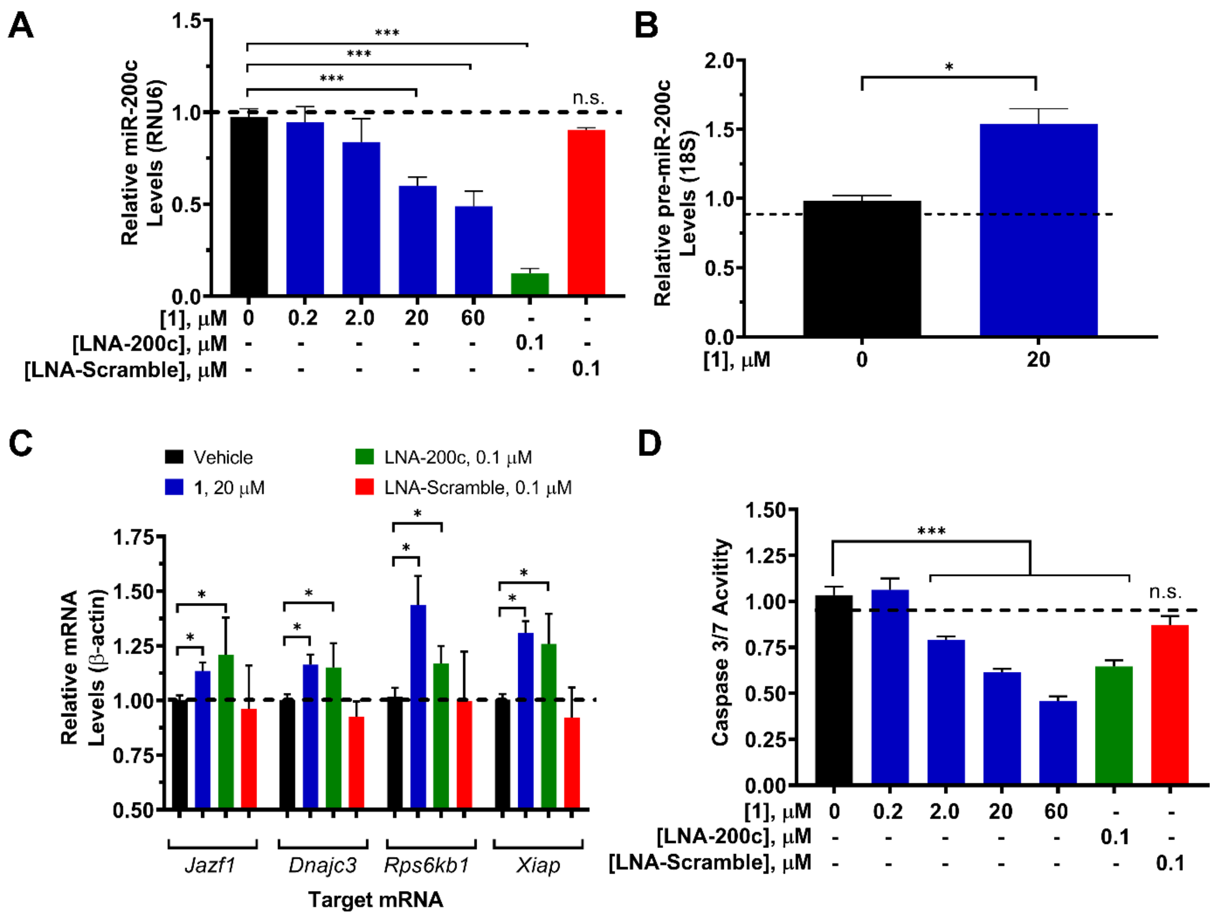

**Figure S5: Compound 1 selectively inhibits miR-200c biogenesis, de-represses downstream targets, and reverses a miR-200c-mediated T2D phenotype.** **A**) Effect of **1** on mature miR-200c levels, as determined by RT-qPCR (n = 5). **B**) Effect of **1** on pre-miR-200c levels, as determined by RT-qPCR (n = 4). **C**) Effect of **1** on miR-200c's direct mRNA targets: *Rps6kb1*, *Dnajc3*, *Xiap* and *Jazf1*, as determined by RT-qPCR (n = 3). **D**) Effect of **1** on Caspase 3/7 shows a 20% reduction in Caspase3/7 activity (n = 8). *, p<0.05; **, p<0.01; ***, p<0.001. All p-values were calculated by a two-tailed Student t-test. All data are reported as mean ± S.E.M.

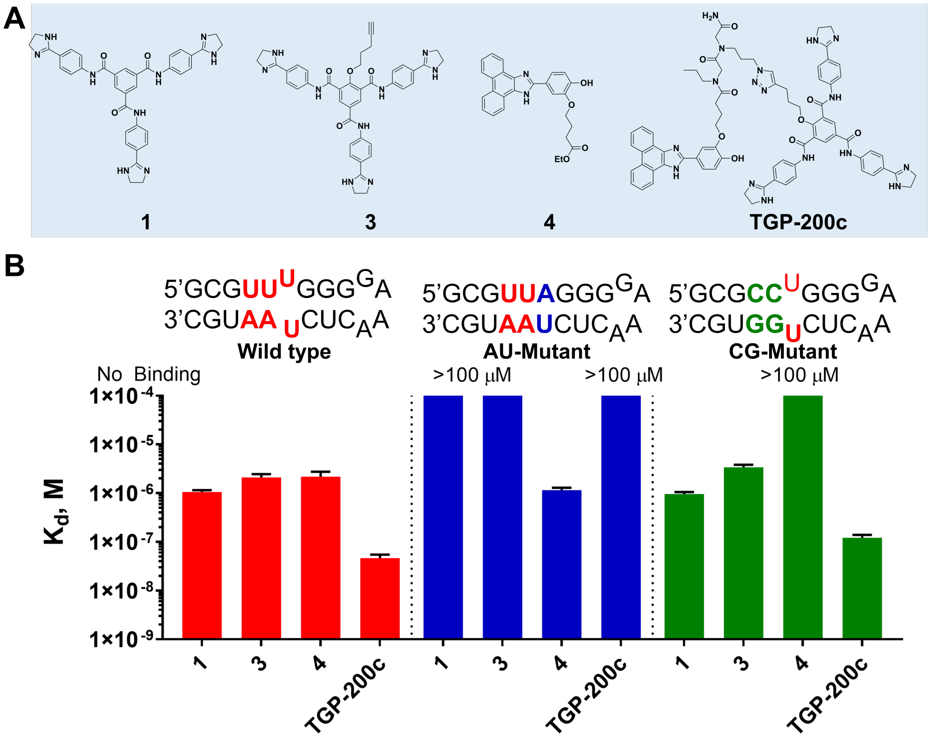

**Figure S6: Small molecules selectively bind to models of miR-200c.** **A**) Structures of small molecules: **3** is a derivative of **1** with an alkyne handle used for modular assembly; **2** binds the UA base pairs adjacent to miR-200c's Dicer processing site; and **4** is an ester derivative of **2**, which selectively binds the UA/UA pairs (2) adjacent to the Dicer site. **TGP-200c** is the optimal dimer with one propylamine spacer separating the **2** and **4** RNA-binding modules. **B**) Summary of $K_d$'s measured for each molecule to: (i) wild type miR-200c's Dicer site; (ii) mutated miR-200c Dicer site with no U/U loop ("AU-mutant"); and (iii) mutated miR-200c Dicer site in which the AU pairs were converted to GC pairs ("GC-mutant") (n =3). Compound **1** bound the wild type RNA, AU-Mutant, and GC-Mutant with $K_d$'s of 1.1 ± 0.1, 1 ± 0.1, and >100 μM respectively. Compound **3** bound similarly to its parent compound **1** with affinities of 2.0 ± 0.1, 3.4 ± 0.4, and > 100 μM, respectively. Compound **4** bound only the wild type RNA and AU-mutant with $K_d$'s of 2.2 ± 0.6 and 1.2 ± 0.1 μM, respectively. No saturable binding was observed for **4** to the GC-mutant RNA. The $K_d$'s for **TGP-200c** and the wild type, AU-mutant, and GC-mutant were 0.050 ± 0.008, >100, and 0.121 ± 0.02 μM, respectively.

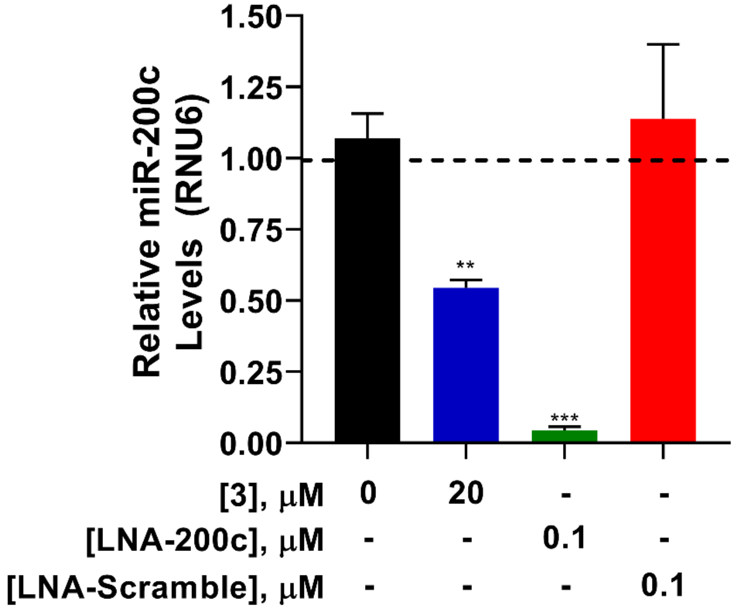

**Figure S7: Chemical probe 3, used for modular assembly, reduces mature miR-200c levels with activity similar to that of parent compound 1.** Compound **3** inhibits miR-200c levels by ~50% at 20 μM, similar to **1** (n = 3). **, p<0.01; ***, p<0.001. All p-values were calculated by a two-tailed Student t-test. All datta are reported as mean ± S.E.M.

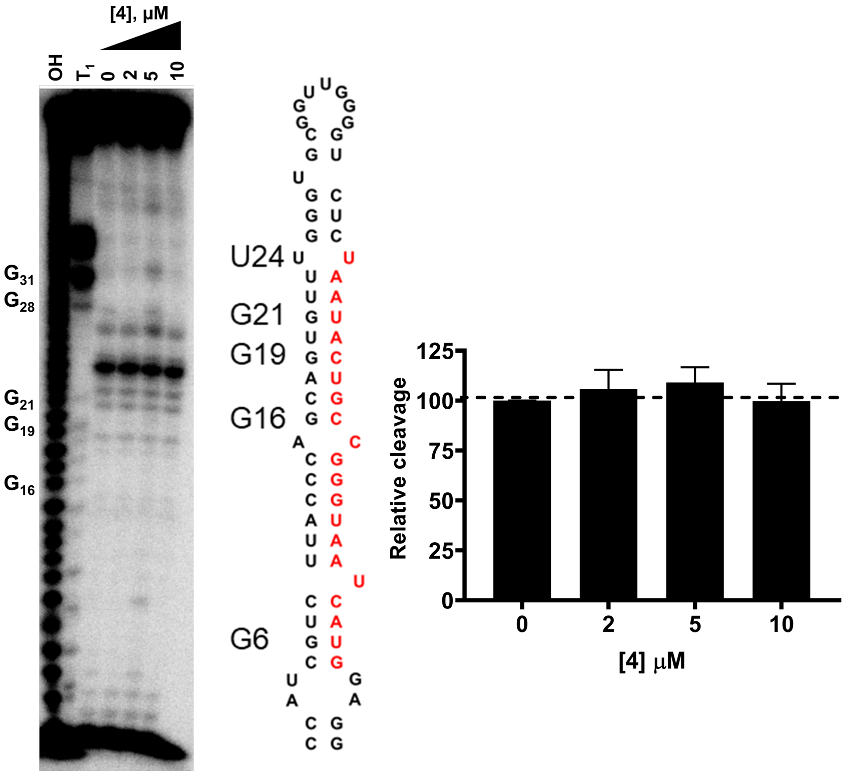

**Figure S8: AU base pair binder 4 does not inhibit the *in vitro* Dicer processing of wild type pre-miR-200c.** All data are reported as the mean ± S.E.M (n = 3).

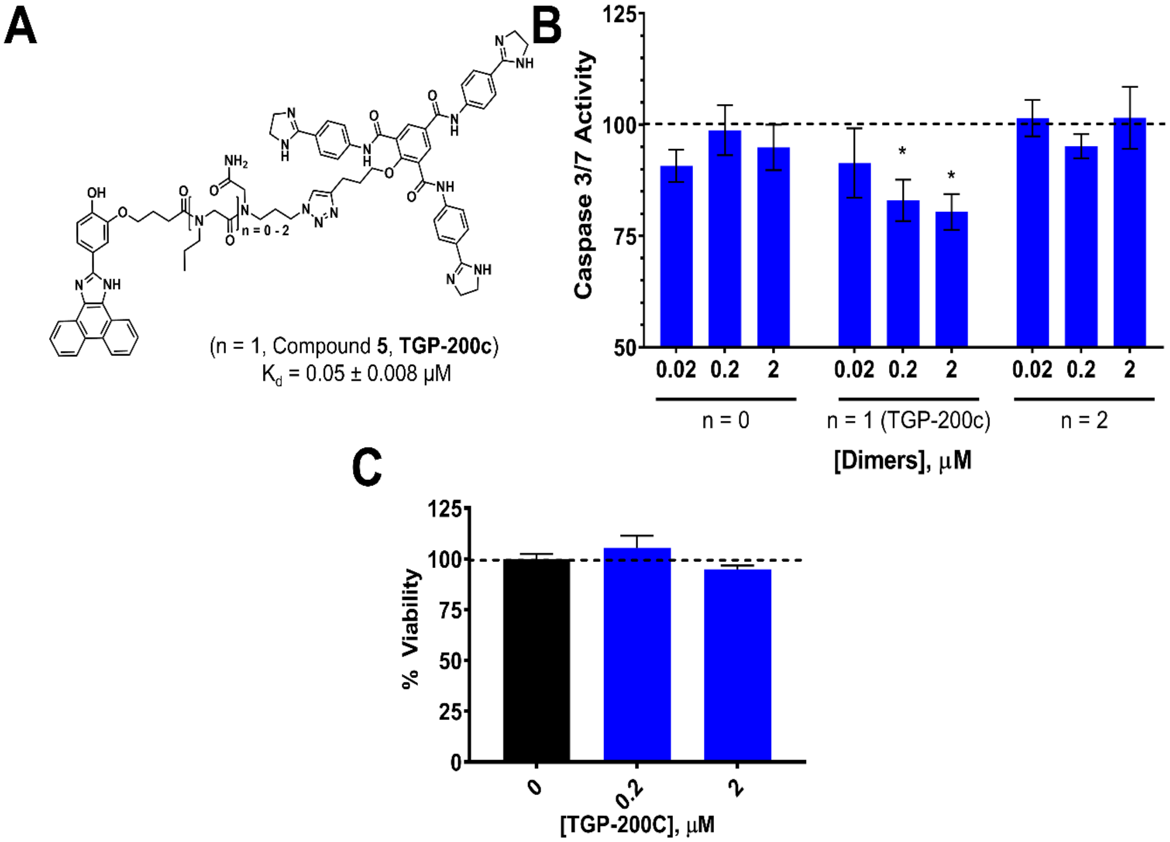

**Figure S9: TGP-200c is the optimal dimer that reduces apoptosis as measured by a Caspase 3/7 assay. A & B)** The library of dimers was screened in MIN6 cells transfected with pre-miR-200c for reducing Caspase 3/7 activity in dose response to identify the optimal dimer, **TGP-200c** (one propylamine spacing unit) (n = 6). **C**) Toxicity of **TGP-200c** as determined by a cell viability assay, demonstrating the compound has no toxicity at the bioactive concentrations (n = 8). *, $p < 0.05$. All p-values were calculated by a two-tailed Student t-test. All data are reported as mean ± S.E.M..

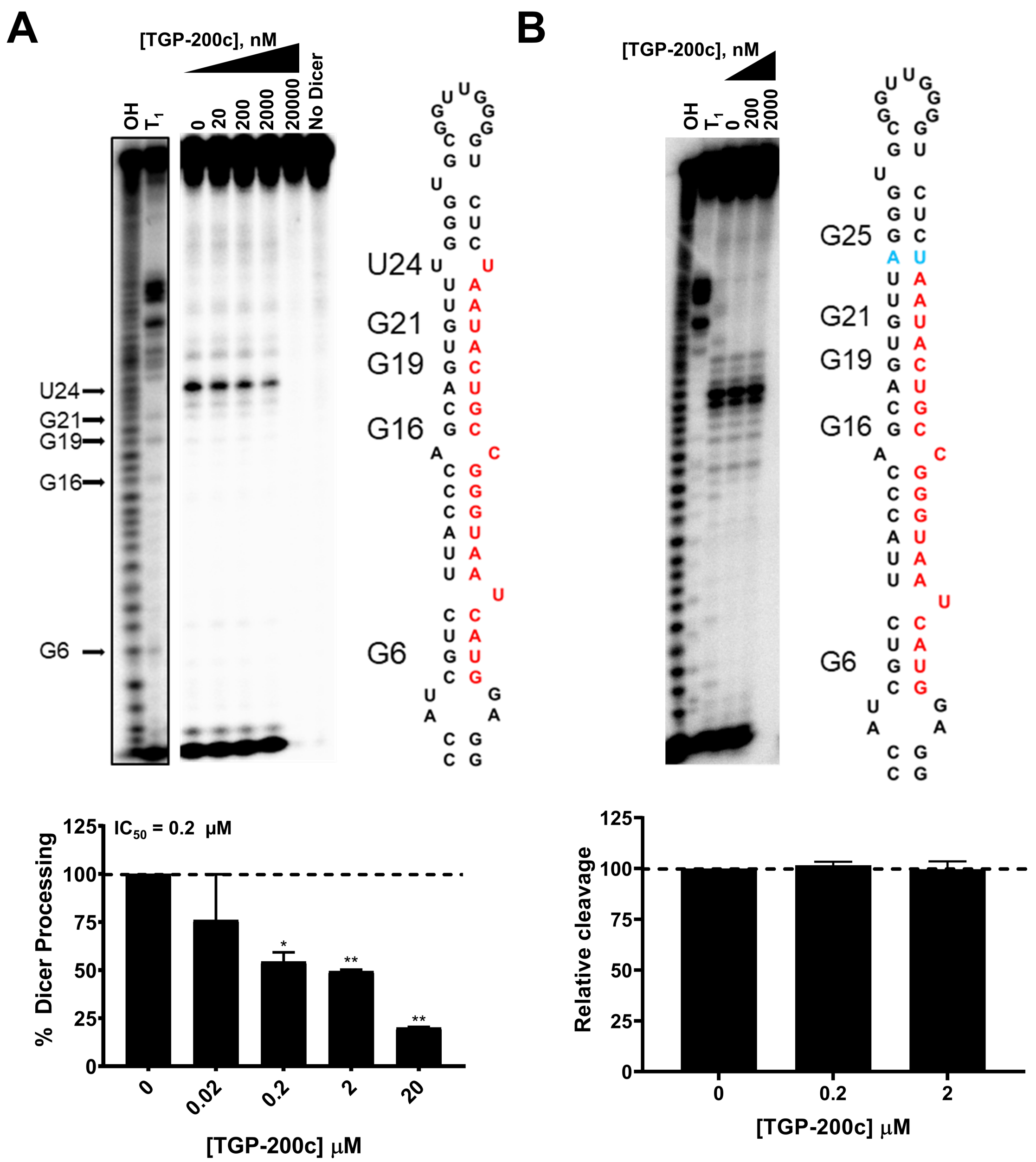

**Figure S10: TGP-200c inhibits the *in vitro* Dicer processing of wild type but not mutant pre-miR-200c.** **A**) Representative gel image of the inhibition of Dicer processing of pre-miR-200c by **TGP-200c** *in vitro,* with an $IC_{50}$ of 200 nM (n = 3). **B**) Representative gel image of the inhibition of Dicer processing of mutant pre-miR-200c by **TGP-200c** *in vitro,* showing no effect (n = 3). *, p < 0.05; **, p < 0.01. All p-values were calculated by a two-tailed Student tt-test. All dat are reported as mean ± S.E.M.

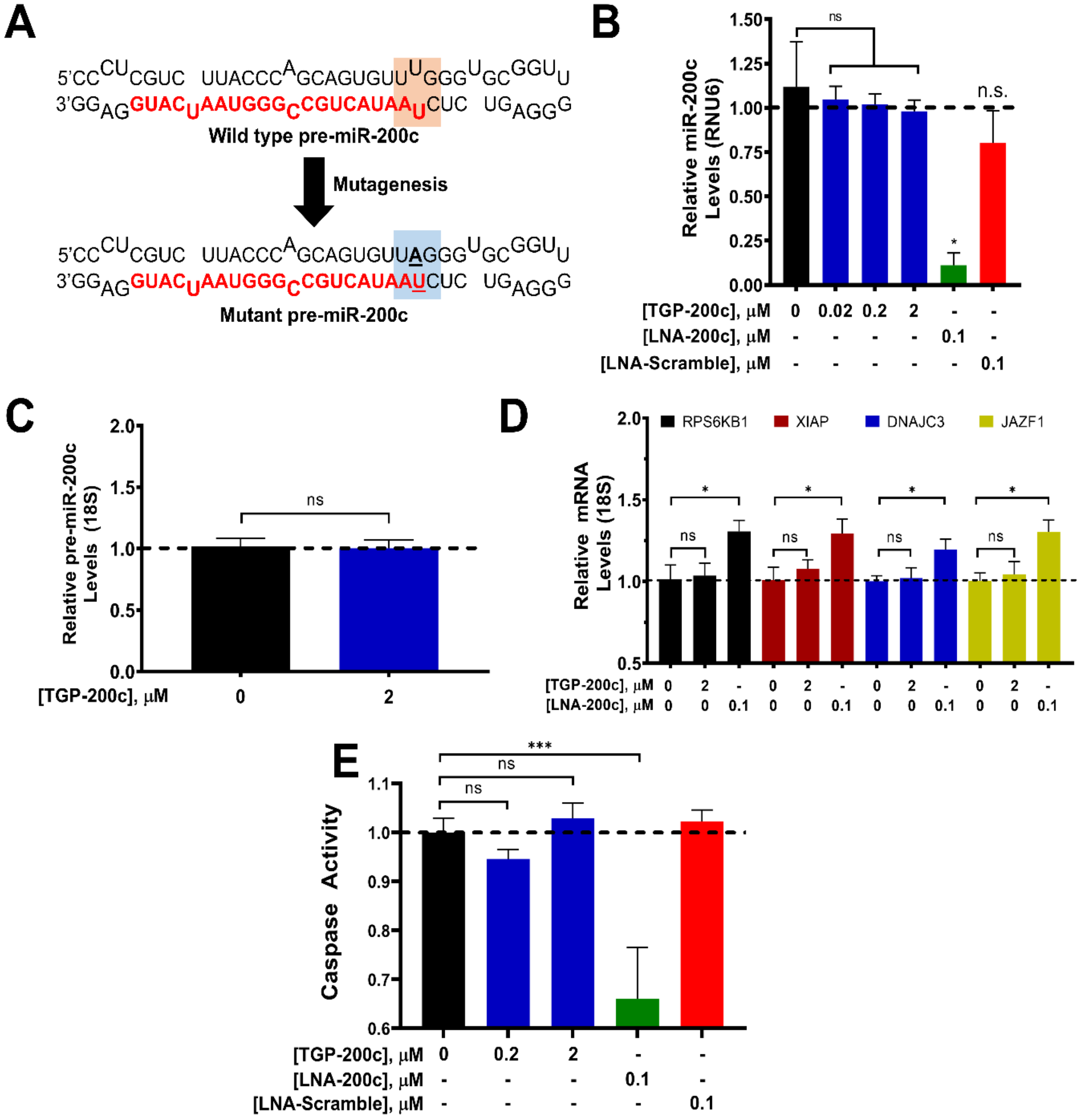

**Figure S11: TGP-200c does not inhibit the biogenesis of mutant pre-miR-200c and its associated downstream targets in MIN6 cells. A**) Secondary structures of WT and mutant pre-miR-200c. In the mutant, the internal loop at the Dicer site is mutated to base pairs, while maintaining the seed sequence of the mature sequence. **B**) Effect of **TGP-200c** on mature miR-200c levels emanating from mutant pre-miR-200c in MIN6 cells, as determined by RT-qPCR (n = 6). Mature miR 200-c expression levels are unchanged by treatment with **TGP-200c**

but reduced by treatment with LNA-200c. **C**) Effect of **TGP-200c** on mutant pre-miR-200c levels, as determined by RT-qPCR (n = 12). **D**) Effect of **TGP-200c** on miR-200c's direct mRNA targets *Rps6kb1*, *Dnajc3*, *Xiap* and *Jazf1* in MIN6 cells expressing mutant pre-miR-200c, which does not bind **TGP-200c**, as determined by RT-qPCR (n = 4). **E**) Effect of **TGP-200c** on Caspase 3/7 activity in MIN6 cells expressing mutant pre-miR-200c (n = 16). *, $p<0.05$; **, $p<0.01$. All p-values were calculated by a two-tailed Student t-test. All data are reported as mean ± S.E.M.

#### Synthetic Methods and Characterization

**Abbreviations:**

AcOH: Acetic acid

DCM: Dichloromethane

DIC: *N*,*N*'-Diisopropylcarbodiimide

DIEA: Diisopropyl ethyl amine

DMF: Dimethylformamide

DMSO: Dimethyl sulfoxide

EDTA: Ethylenediaminetetraacetic acid

$Et_2O$: Diethyl ether

EtOH: Ethanol

HATU: Hexafluorophosphate azabenzotriazole tetremethyl uronium

HOAt: 1-hydroxy-7-azabenzotriazole

HPLC: High-Performance Liquid Chromatography

MeOH: Methanol

TEA: Triethylamine

TFA: Trifluoroacetic Acid

THF: Tetrahydrofuran

**General Methods:** HPLC purification was completed using a Waters 1525 Binary HPLC Pump equipped with a Waters 2487 Dual Absorbance Detector system and a reverse phase Atlantis® Prep T3 C18 5 µM column. The gradient used for purification was from 0% to 100% solvent B (100% MeOH + 0.1% TFA) in solvent A ($H_2O$ + 0.1% TFA) over 100 min. Purity was evaluated with an analytical HPLC equipped with a reverse phase Waters Symmetry C18 5 µm 4.6 × 150 mm column with a flow rate of 1 mL/min from 0% to 100% solvent B in solvent A over 60 min. Absorbance was detected at 220 nm and 254 nm.

Mass spectra were recorded on a 4800 plus MALDI TOF/TOF analyzer. NMR spectra were measured by using a Bruker 400 UltraShield™. Chemical shifts are shown in ppm relative to TMS. Coupling constants (*J*) are reported in Hertz.

Chemicals were purchased from commercial sources and used without further purification as follows: Rink Resin SS (loading = 0.50 mmol/g). *N,N'*-diisopropylcarbodiimide (DIC), 9,10-Phenanthrenequinone from Chem-Impex Int'l Inc.; 1.3-dimethyluracil, Dimethyl 1,3-acetonedicarboxylate, 5-Chloro-1-pentyne, 4-aminobenzonitrile, 3,4-dihydroxybenzaldehyde, 4-bromobutyric acid ethyl ester were obtained from Combi-Blocks; HATU and trifluoroacetic acid from Oakwood Chemical; 1-propylamine, acetyl chloride, benzyl bromide and Phosphorus(V) oxychloride from Alfa Aesar; 2-bromoacetic acid, *N*, *N*-diisopropylethyl amine, and propargylamine from Sigma Aldrich; Oxalyl chloride and ethylenediamine from Acros Organics; and dimethyl sulfoxide (DMSO, anhydrous) and *N*,*N*-dimethylformamide (DMF, anhydrous) from EMD.

#### Synthetic Schemes and Methods for Small Molecules

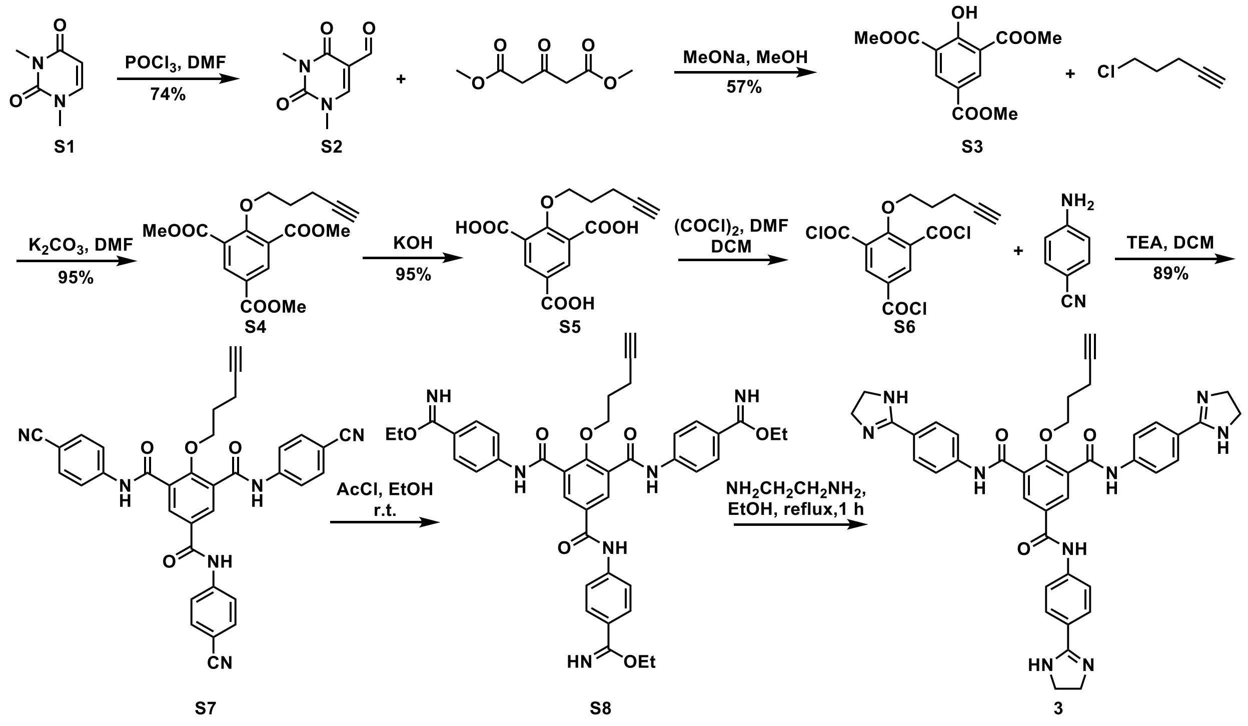

**Scheme 1**: Synthesis of compound **3**

**Synthesis of S2.** **S2** was synthesized by modification of route reported in reference (3). To a solution of phosphorus oxychloride (11.5 g, 0.075 mol) in 40 mL of DMF at 0 °C was added 1,3-dimethylracil (**S1**) (7 g, 0.05 mol). The reaction mixture was heated at 85 °C for 1 h. Then, the reaction was cooled to room temperature and concentrated *in vacuo*. The residue was extracted with ethyl acetate from $H_2O$ three times, and the combined organic layers were dried over *anhyd.* $Na_2SO_4$ and concentrated *in vacuo*. The residue was purified by column chromatography (240 mesh silica gel) to give the 5-formyl-1,3-dimethyluracil (**S2**) as a white solid (6.2 g, 0.044 mol, 74%). **$^1$H NMR** (400 MHz, $CDCl_3$) δ (ppm) 10.04 (s, 1H), 8.06 (s, 1H), 3.53 (s, 3H), 3.39 (s, 3H); **$^{13}$C NMR** ( 100 MHz, $CDCl_3$) δ (ppm) 186.8, 161.9, 150.9, 147.7, 110.1, 38.0, 27.8; **HRMS** (m/z): calculated for $C_7H_9N_2O_3$ $[M+H]^+$: 169.0608, found: 169.0600.

**Synthesis of S3.** **S3** was synthesized by modification of route reported in reference (3). To a solution of 5-formyl-1,3-dimethyluracil (**S2**) (2g, 11.9 mmol) and dimethyl-1,3-acetonedicarboxylate (8.29 g, 47.6 mmol) was added MeONa (2.12 g, 39.3 mmol). The reaction mixture was heated under reflux for 3 h and then cooled to room temperatturer. The precipitate was collected by filtration and washed with $H_2O$. Then, the solid was acidified with 3 M HCl and extracted with ethyl acetate from $H_2O$ to give the dimethyl 5-acetoxy-4-hydroxyisophthalate (**S3**) as a white solid (1.81 g, 6.75 mmol, 57%). **$^1$H NMR** (400 MHz, $CDCl_3$) δ (ppm) 12.26 (s, 1H), 8.72 (s, 2H), 3.99 (s, 6H), 3.93 (s, 3H); **$^{13}$C NMR** ( 100 MHz, $CDCl_3$) δ (ppm) 167.3, 165.2, 164.6, 137.5, 120.7, 116.6, 52.7, 52.3; **HRMS** (m/z): calculated for $C_{12}H_{13}O_7$ $[M+H]^+$: 269.0656, found: 269.0636.

**Synthesis of S4.** To a solution of **S3** (670 mg, 2.5 mmol), 5-chloro-1-pentyne (510 mg, 5 mmol) and KI (830 mg, 5 mmol) in 5 mL DMF was added $K_2CO_3$ (690 mg, 5 mmol). The reaction mixture was stirred at 90 °C overnight. Then, the reaction mixture was diluted with $H_2O$ and extracted with ethyl acetate three times. The combined organic layers were washed with brine, dried over *anhyd.* $Na_2SO_4$ and concentrated *in vacuo*. The residue was purified by column chromatography to give the dimethyl 5-acetoxy-4-(pent-4-yn-1-yloxy)isophthalate (**S4**) as a white solid (798 mg, 2.39 mmol, 95%). **$^1$H NMR** (400 MHz, $CDCl_3$) δ (ppm) 8.54 (s, 2H), 4.14 (t, *J*= 6 Hz, 2H); 3.93 (s, 6H), 3.92 (s, 3H), 2.43 (td, *J*=7.1, 2.6 Hz, 2H), 2.03 (m, 2H), 1.95 (t, *J*= 2.6 Hz, 1 H); **$^{13}$C NMR** ( 100 MHz, $CDCl_3$) δ (ppm), 165.4, 165.0, 161.7, 136.0, 126.7, 125.3, 83.6, 75.1, 68.7, 52.6, 52.5, 29.0, 15.0; **HRMS** (m/z): calculated for $C_{17}H_{19}O_7$ $[M+H]^+$: 335.1125, found: 335.1154.

**Synthesis of S5.** To a solution of **S4** (798 mg, 2.39 mmol) in 10 mL of THF/$H_2O$ (8:2) was added KOH (669 mg, 11.9 mmol). The reaction mixture was stirred at 50 °C overnight. The reaction mixture was acidified with 3M HCl, and the precipitate was collected by filtration to give the 2-(pent-4-yn-1-yloxy)benzene-1,3,5-tricarboxylic acid (**S5**) as a white solid (660 mg, 2.26 mmol, 95%). **$^1$H NMR** (400 MHz, $CDCl_3$) δ (ppm) 13.38 (br, 3H), 8.30 (s, 2H), 4.09 (t, *J*=6.2 Hz,

2H), 2.77(t, *J*=2.6 Hz, 1H), 2.31(td, *J*=7.3, 2.6 Hz, 2H), 1.87(m, 2H); $^{13}$**C NMR** ( 100 MHz, $CDCl_3$) δ (ppm) 166.4, 165.7, 160.2, 134.3, 127.7, 125.5, 83.9, 74.6, 71.4, 28.8, 14.5; **HRMS** (m/z): calculated for $C_9H_7O_7$ $[M+H]^+$: 227.0186, found: 227.0177.

**Synthesis of S7.** To a suspension of **S5** (200 mg, 0.684 mmol) and oxalyl chloride (522 mg, 4.11 mmol) in 5 mL of dry DCM was added 2 drops of dry DMF. The reaction mixture was stirred at room temperature for 3 h and then concentrated *in vacuo*. The residue was re-suspended in 5 mL of dry THF. To this solution was added 4-aminobenzonitrile (485 mg, 4.11 mmol) and TEA (691 mg, 6.84 mmol). The reaction mixture was stirred at room temperature for 3h. This reaction mixture was diluted with ethyl acetate and washed with 3 M HCl three times and then with 3 M KOH three times. The organic phase was then washed with brine, dried over *anhyd.* $Na_2SO_4$, and concentrated *in vacuo* to give the pure product **S7** as a white solid (350 mg, 0.59 mmol, 87%). $^1$**H NMR** (400 MHz, DMSO-*d*$^6$) δ (ppm) 11.01(s, 2H), 11.76 (s, 1H), 8.33 (s, 2H), 8.79 (d, *J*=8.8 Hz, 2H), 7.94 (d, *J*=8.8 Hz, 4H), 7.85 (m, 6H), 4.14(t, *J*=6.0 Hz, 2H), 2.76(t, *J*=2.6 Hz, 1H), 2.15 (td, *J*=7.3, 2.7 Hz, 2H), 1.75(m, 2H); $^{13}$**C NMR** ( 100 MHz, DMSO-*d*$^6$) δ (ppm) 165.0, 164.5, 156.6, 143.7, 143.4, 133.3, 133.1, 130.9, 130.3, 128.6, 120.2, 119.9, 119.1, 105.5, 105.3, 83.3, 73.7, 71.4, 28.6, 14.3; **HRMS** (m/z): calculated for $C_{35}H_{25}N_6O_4$ $[M+H]^+$: 593.1932, found: 593.1883.

**Synthesis of 3.** To a solution of **S7** (200 mg, 0.338 mmol) in 6 mL of EtOH was added 3.5 mL of AcCl. The reaction mixture was stirred at room temperature for 3 d. Then, the reaction mixture was bubbled with air and then $Et_2O$ was added. The precipitate was collected by filtration and dried *in vacuo* to afford **S8**, which was used directly for the next step. The residue was resuspended in EtOH, followed by the addition of 0.5 mL of ethylenediamine. The mixture was stirred under reflux for 2 h. Then, the reaction mixture was cooled down to room temperature and concentrated *in vacuo*. The residue was purified by HPLC to give **3** as a TFA salt. $^1$**H NMR** (400 MHz, $CD_3OD$) δ (ppm) 8.44(s, 2H), 8.06 (m, 6H), 7.90 (m, 6H), 4.25 (t, *J*=5.9 Hz, 2H), 2.24 (td, *J*=7.2, 2.7 Hz, 2H), 2.00 (t, *J*=2.6 Hz, 1H), 1.87 (m, 2H); $^{13}$**C NMR** (100 MHz, $CD_3OD$) δ (ppm) 167.1, 166.7, 166.3, 158.5, 146.1, 145.7, 132.7, 131.9, 131.2, 130.7, 130.5, 121.7, 121.2, 118.8, 118.5, 83.5, 76.1, 70.3, 45.9, 37.9, 30.2, 15.7;**HRMS** (m/z): calculated for $C_{41}H_{40}N_9O_4$ $[M+H]^+$: 722.3198, found: 722.3192.

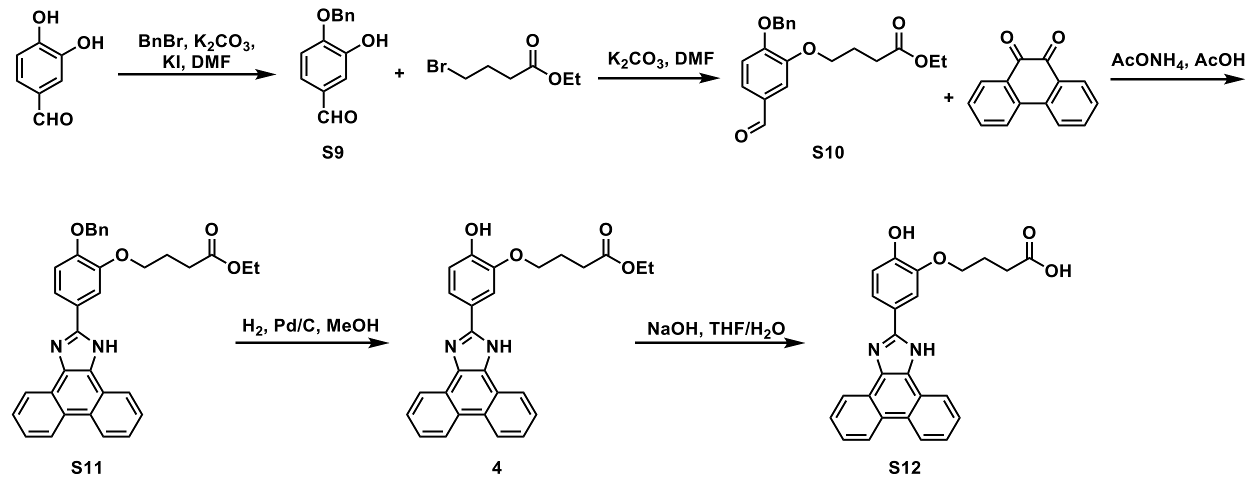

**Scheme 2**: Synthesis of **S12**.

**Synthesis of S10.** To a solution of 4-benzyloxy)-3-hydroxybenzaldehyde (**S9**; 456 mg, 2 mmol) (4) and ethyl 4-bromobutyrate (466 mg, 2.4 mmol) in 5 mL of DMF was added $K_2CO_3$ (331 mg, 2.4 mmol). The reaction mixture was stirred at room temperature overnight, and then diluted with $H_2O$ and extracted with ethyl acetate three times. The organic phase was washed with brine, dried over *anhyd.* $Na_2SO_4$, and concentrated *in vacuo*. The residue was purified by column chromatography to give **S10** as a colorless oil (500 mg, 1.46 mmol, 73%). **$^1$H NMR** (400 MHz, $CDCl_3$) δ (ppm) 9.83(s, 1H), 7.31-7.46(m, 7H), 6.99(d, *J*= 8.2 Hz, 1H), 5.22 (s, 2H), 4.11-4.17(m, 4H), 2.55(t, *J*=7.4 Hz, 2H), 2.17-2.21 (m, 2H), 1.26(t, *J*=7.2 Hz, 3H); **$^{13}$C NMR** (100 MHz, $CDCl_3$) δ (ppm) 190.9, 173.1, 154.0, 149.3, 136.2, 130.3, 128.6, 128.1, 127.0, 126.5, 112.8, 111.2, 70.7, 67.9, 60.5, 30.7, 24.5, 14.2; **HRMS** (m/z): calculated for $C_{20}H_{23}O_5$ $[M+H]^+$: 343.1540, found: 343.1513.

**Synthesis of S11.** A solution of **S10** (342 mg, 1 mmol), phenanthrenequinone (208 mg, 1 mmol) and $AcONH_4$(308 mg, 4 mmol) in 4 mL of AcOH was stirred at reflux overnight. Then the reaction mixture was cooled to RT and to this solution was added 10 mL of $H_2O$. The solid was extracted by ethyl acetate from $H_2O$. The organic phase was washed with brine, dried over *anhyd.* $Na_2SO_4$ and concentrated *in vacuo*. The residue was washed with $Et_2O$ to give **S11** as a pale white solid (400 mg, 0.754 mmol, 75%). **$^1$H NMR** (TFA salt, 400 MHz, DMSO-$d^6$) δ (ppm) 8.93 (d, $J$=8.3 Hz, 2H), 8.59(d, $J$=7.5 Hz, 2H), 7.89 (m, 2H), 7.79-7.85(m, 2H), 7.70-7.76(m, 2H), 7.50-7.54(m, 2H), 7.33-7.46 (m, 4H), 5.27 (s, 2H), 4.21 (t, $J$=6.2 Hz, 2H), 4.08 (q, $J$=7.1 Hz, 2H), 2.56(t, $J$=7.3 Hz, 2H), 2.05-2.10 (m, 2H), 1.18 (t, $J$=7.1 Hz, 3H); **$^{13}$C NMR** (TFA salt, 100 MHz, DMSO-$d^6$) δ (ppm) 172.6, 149.2, 148.7, 137.1, 128.5, 127.9, 127.6, 127.5, 127.1, 125.1, 125.0, 124.1, 123.8, 123.6, 122.4, 121.9, 119.4, 114.4, 111.7, 70.0, 67.8, 59.9, 30.1, 24.5, 14.1; **HRMS** (m/z): calculated for $C_{34}H_{31}N_2O_4$ $[M+H]^+$: 531.2278, found: 531.2203.

**Synthesis of 4 and S12.** To a solution of **S11** (100 mg, 0.189 mmol) in MeOH was added 10% Pd/C (10 mg) and the reaction mixture was shaken at room temperature under an atmosphere of $H_2$ for 3 h. The Pd/C was filtered off through Celite, and the filtrate was concentrated *in vacuo* affording crude **4** which was used directly for next step without further purification. The residue was dissolved in 2 mL of THF, and to this solution was added a solution of NaOH (22.6 mg, 0.566 mmol) in 1 mL of $H_2O$. The mixture was stirred at 50 °C for 3 h. The reaction mixture was acidified with 3M HCl to pH=7-8. The precipitate was collected by filtration to give **S12** as a pale white solid (55 mg, 0.133 mmol, 70%). **$^1$H NMR** (400 MHz, DMSO-$d^6$) δ (ppm) 10.1 (br, 1H), 8.98 (d, $J$=8.3 Hz, 2H), 8.83 (d, $J$=7.8 Hz, 2H), 8.12 (s, 1H), 7.95 (dd, $J$=8.3, 1.9 Hz, 1H), 7.85-7.88(m, 2H), 7.77-7.81(m, 2H), 7.13 (d, $J$=8.4 Hz, 1H), 4.23 (t, $J$=6.8 Hz, 2H), 2.53 (t, $J$=7.3 Hz, 2H), 2.02-2.09 (m, 2H); **$^{13}$C NMR** (100 MHz, DMSO-$d^6$) δ (ppm) 174.3, 151.2, 147.6, 147.2, 128.4, 127.9, 127.3, 126.5, 124.2, 123.1, 122.4, 121.0, 116.1, 113.4, 630.1, 681, 30.1, 24.3; **HRMS** (m/z): calculated for $C_{25}H_{21}N_2O_4$ $[M+H]^+$: 413.1496, found: 413.1450.

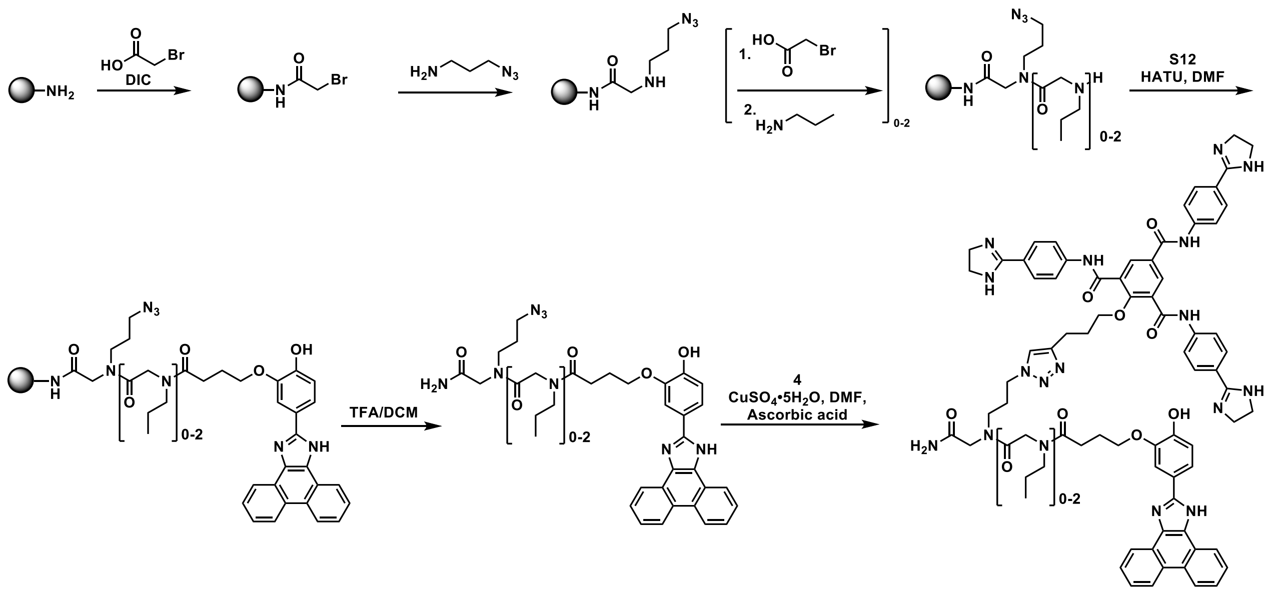

**Scheme 3**: Synthesis of a library of dimers targeting pre-miR-200c.

**Deprotection of peptoid.** Fmoc-protected rink resin (400 mg, 0.2 mmol) in 5 mL of 20% piperidine in DMF was shaken at room temperature for 30 min. The resin beads were then washed with 5 mL of DMF three times.

**Amide coupling with bromoacetic acid**: The resin was then suspended in 3 mL of DMF followed by the addition of bromoacetic acid (200 mg, 1.44 mmol) and DIC (400 mg, 3.18 mmol). The reaction mixture was shaken at room temperature for 2 h and then washed with 5 mL of DMF three times.

**SN2 substitution with 1-propylamine**: The resin was suspended in 3 mL of DMF followed by the addition of 3-azido-1-propanamine (100 mg, 1 mmol). The mixture was shaken at room temperature for 2 h and then washed with 5 mL of DMF three times.

**Amide coupling with 3**: The **Amide coupling with bromoacetic acid** and **SN2 substitution with 1-propylamine** steps were repeated for 0-2 times. The resin was then suspended in 3 mL of DMF followed by the addition of **S12** (124 mg, 0.3 mmol) and HATU (152 mg, 0.4 mmol). The reaction mixture was shaken at room temperature for 2 h and then washed with 5 mL of DMF three times.

**Cleaving the peptoid off the beads**: The peptoid was cleaved from the beads by shaking in 5 mL of 30% TFA in DCM for 30 min. The solvent was removed in vacuum and purified by HPLC.

**Synthesis of Dimers**: A solution of the peptoid obtained above (1 equiv.), **3** (1 equiv.), $CuSO_4$•5 $H_2O$ (1 equiv.), and ascorbic acid (1 equiv.) in DMF was stirred at room temperature overnight. The product was purified via preparative HPLC

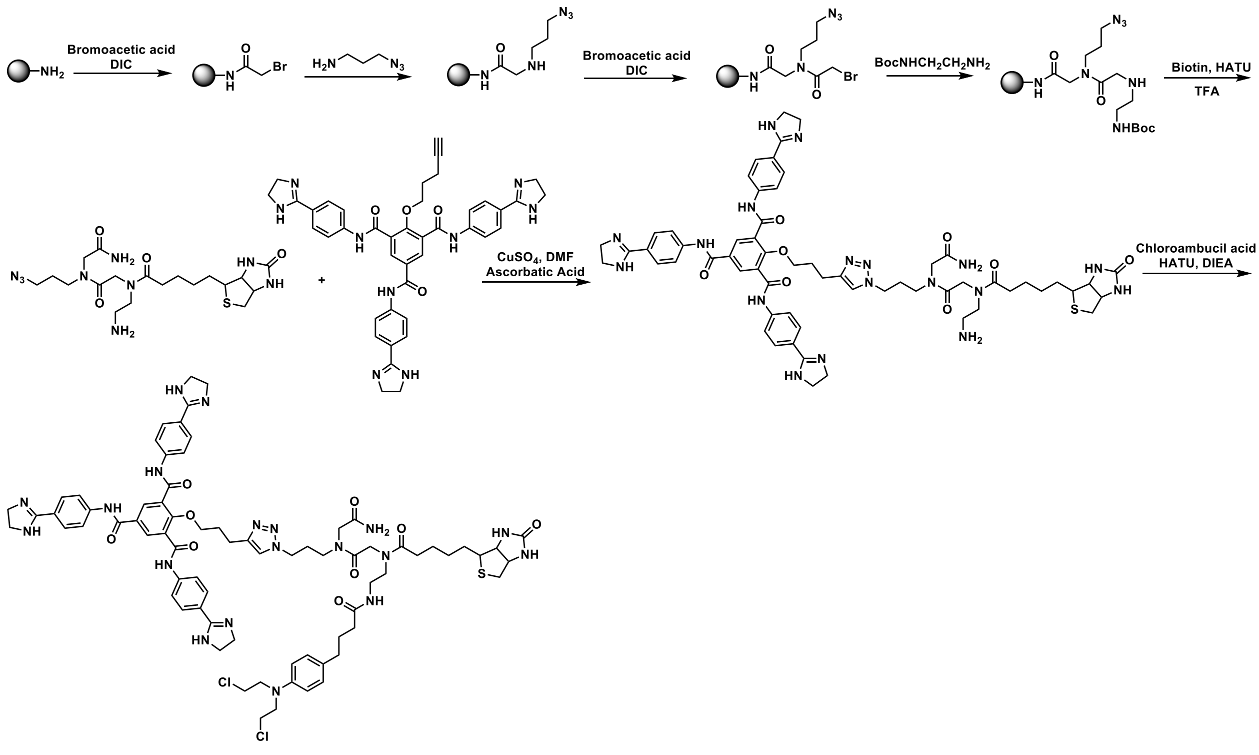

**Scheme 4**. Synthesis of **1-CA-Biotin** (**6**).

**Deprotection of peptoid.** Fmoc-protected rink resin (200 mg, 0.1 mmol) in 5 mL of 20% piperidine in DMF was shaken at room temperature for 30 min. The resin beads were then washed with 5 mL of DMF three times.

**Amide coupling with bromoacetic acid**: The resin was then suspended in 3 mL of DMF followed by the addition of bromoacetic acid (100 mg, 0.72 mmol) and DIC (200 mg, 1.59 mmol). The reaction mixture was shaken at room temperature for 2 h and then washed with 5 mL of DMF three times.

**SN2 substitution with 3-azido-1-propanamine**: The resin was suspended in 3 mL of DMF followed by the addition of 3-azido-1-propanamine (100 mg, 1 mmol). The mixture was shaken at room temperature for 2 h and then washed with 5 mL of DMF three times.

**Amide coupling with bromoacetic acid**: The resin was then suspended in 3 mL of DMF followed by the addition of bromoacetic acid (100 mg, 0.72 mmol) and DIC (200 mg, 1.59 mmol). The reaction mixture was shaken at room temperature for 2 h and then washed with 5 mL of DMF three times.

**SN2 substitution with 1-Boc-ethylenediamine**: The resin was suspended in 3 mL of DMF followed by the addition of N-Boc-ethylenediamine (160 mg, 1 mmol). The mixture was shaken at room temperature for 2 h and then washed with 5 mL of DMF three times.

**Amide coupling with Biotin**: The resin was then suspended in 3 mL of DMSO followed by the addition of Biotin (244 mg, 1 mmol) and HATU (380 mg, 1 mmol). The reaction mixture was shaken at room temperature for 2 h and then washed with 5 mL of DMF three times and 5 mL of DCM three times.

**Cleaving the peptoid off the beads**: The peptoid was cleaved from the beads by shaking in 3 mL of 30% TFA in DCM for 30 min. The solvent was removed under vacuum, and the product was used without further purification.

**1-CA-Biotin**: A solution of the peptoid obtained above (2 mg, 0.004mmol), **3** (2.9 mg, 0.004 mmol), $CuSO_4 \cdot 5H_2O$ (1.0 mg, 0.004 mmol), and ascorbic acid (0.7 mg, 0.004 mmol) in 0.5 mL of DMF was stirred at room temperature overnight. The product was purified by HPLC. The click product was then added to a solution of chlorambucil acid (1.2 mg, 0.004 mmol) and HATU (1.5 mg, 0.004 mmol) in 0.3 mL of DMF, followed by the addition of DIEA (0.782 mg, 0.6 mmol). The reaction mixture was stirred at room temperature for 20 min followed by HPLC purification to give the Chem-CLIP probe **1-CA-Biotin** or **6** (0.6 mg, 0.0004 mmol, 10%)

#### COMPOUND CHARACTERIZATION

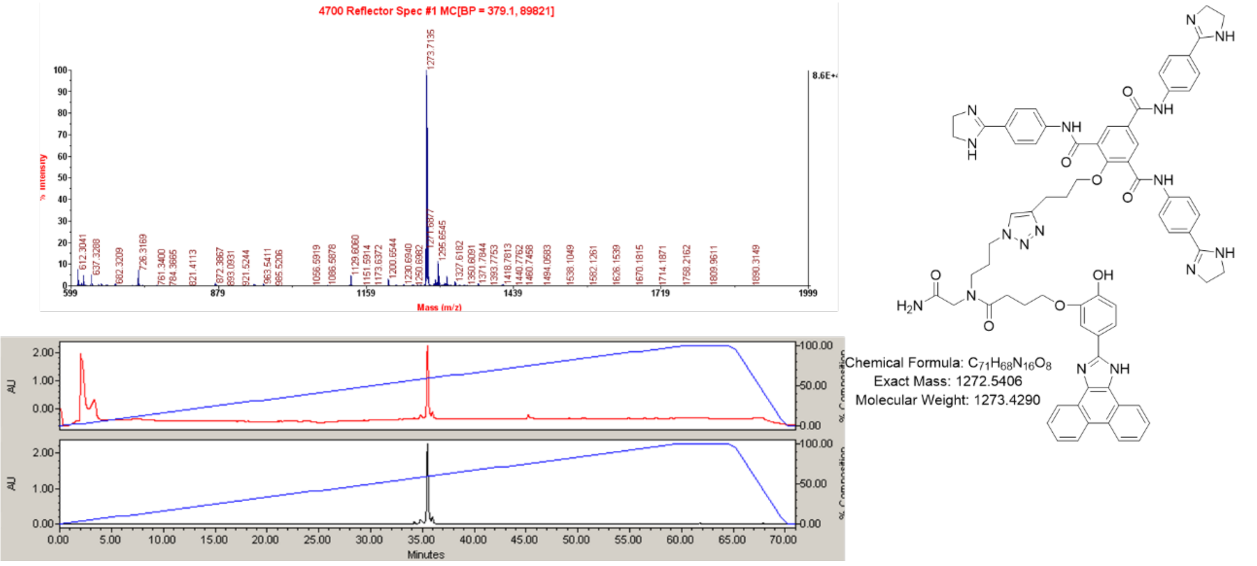

**Figure S11.** Characterization of 200c dimer n = 0.

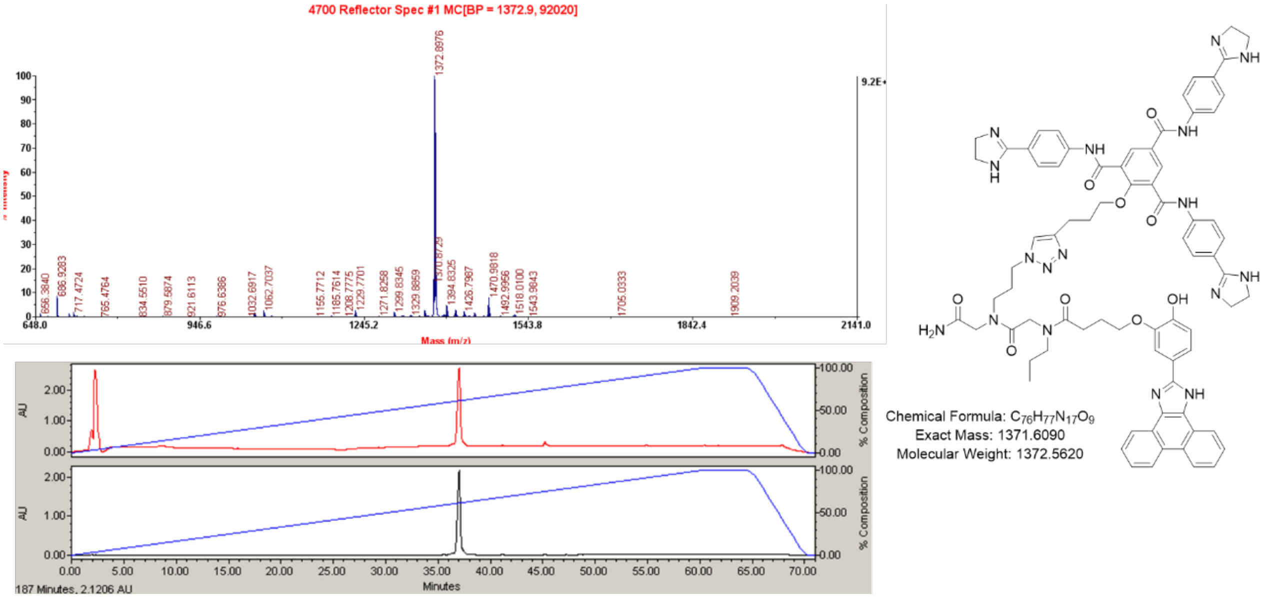

**Figure S12.** Characterization of 200c dimer n = 1.

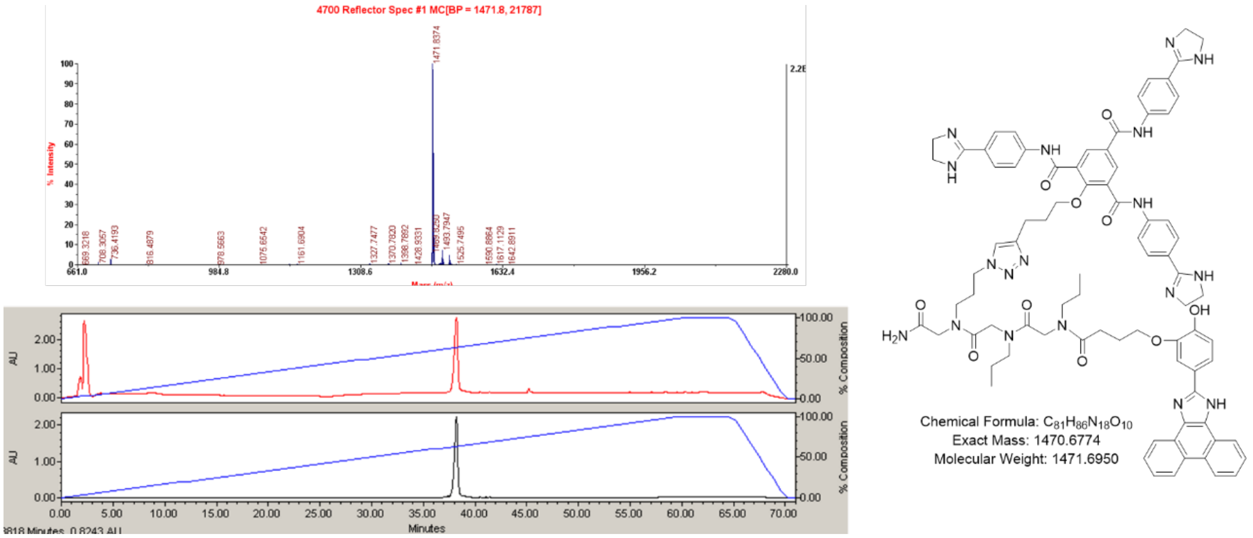

**Figure S13.** Characterization of 200c dimer n = 2.

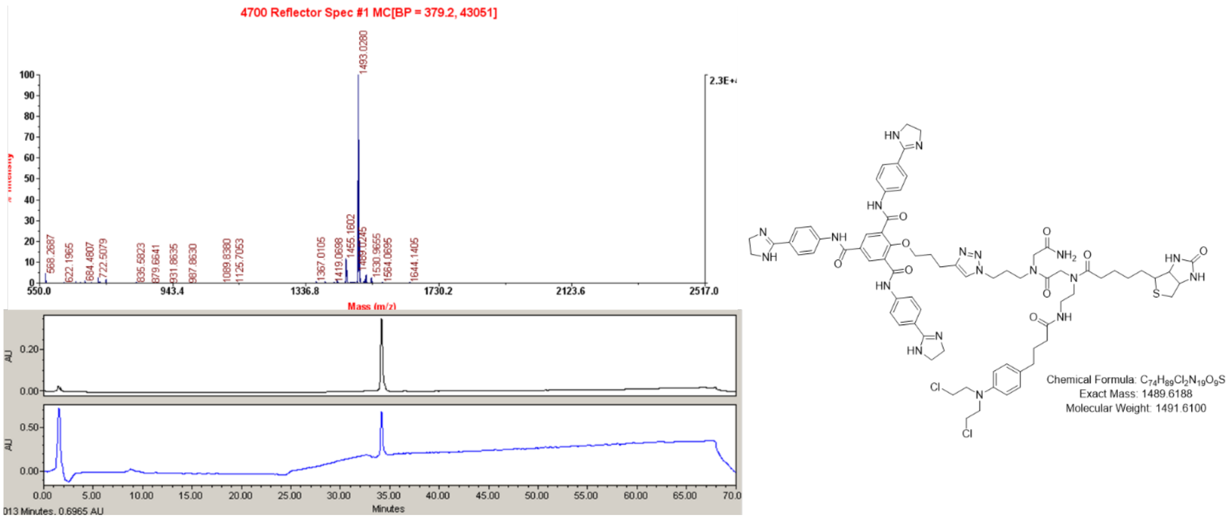

**Figure S14: Characterization of 1-CA-Biotin probe 6.**

### NMR Spectra

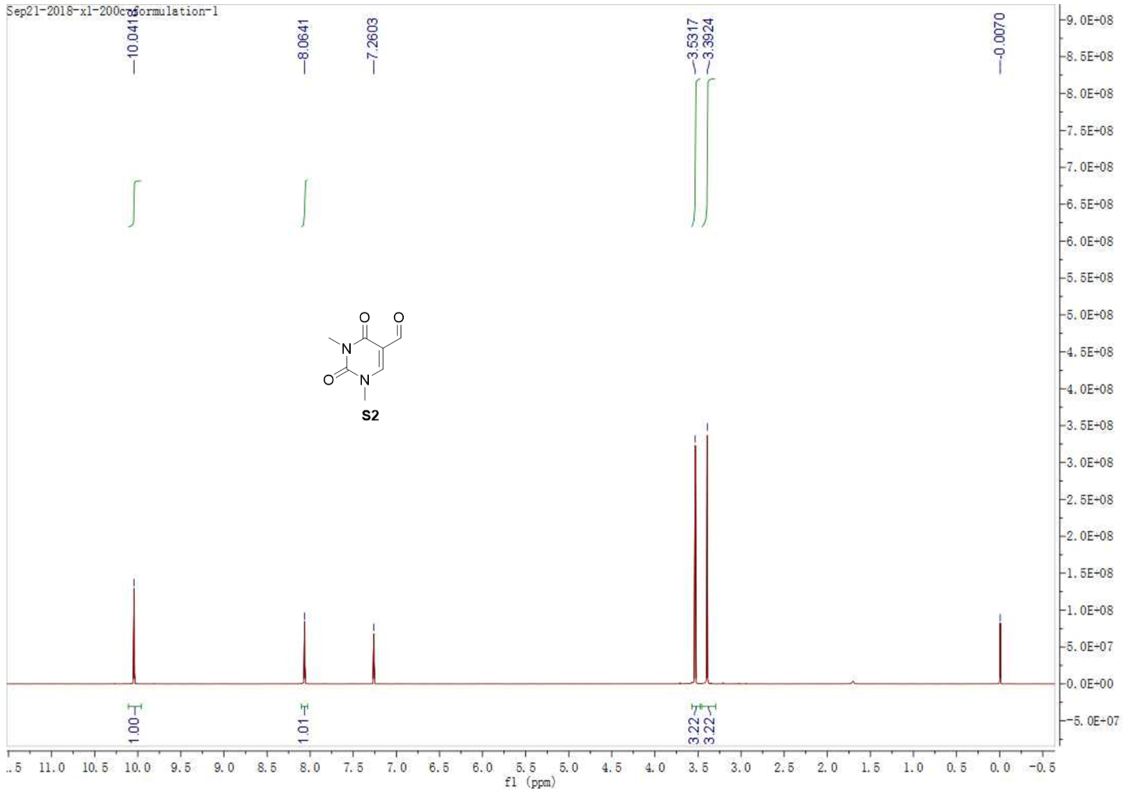
Sep21-2018-xl-200coformulation-1
10.0413
8.0641
7.2603
3.5317
3.3924
-0.0070
S2
1.00
1.01
3.22
3.22
f1 (ppm)

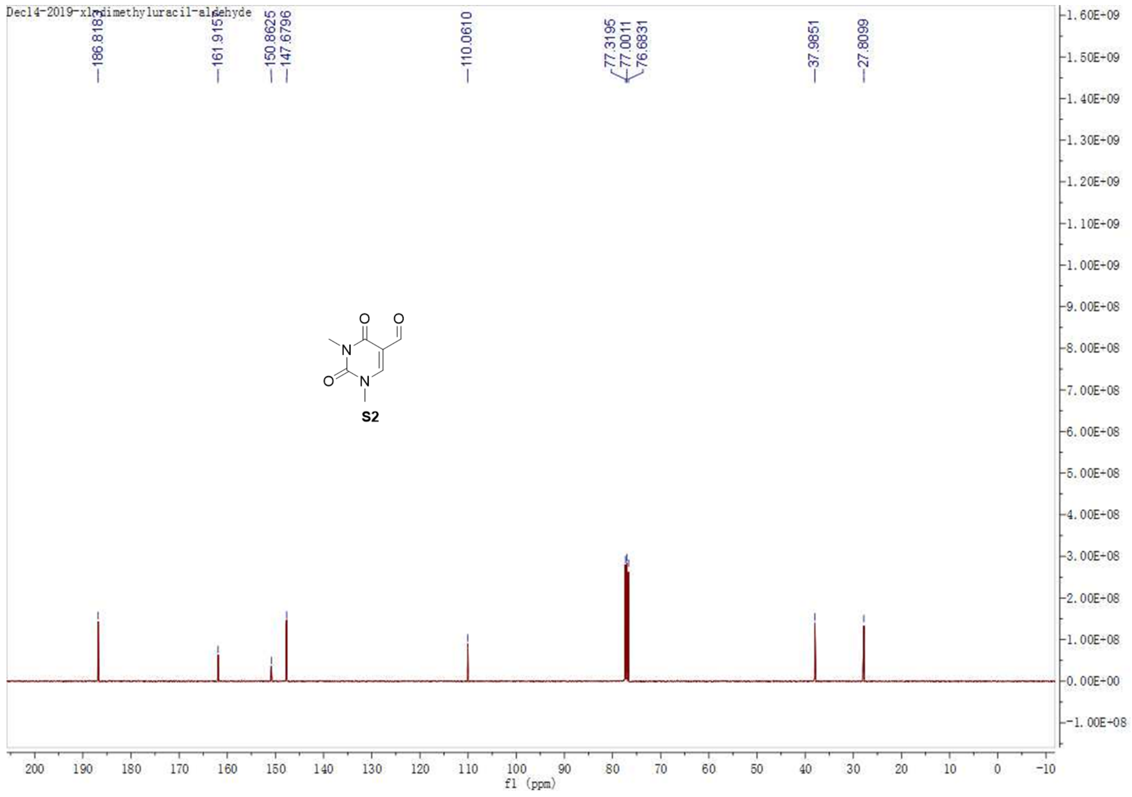
Dec14-2019-xl-dimethyluracil-aldehyde
186.8187
161.9157
150.8625
147.6796
110.0610
77.3195
77.0011
76.6831
37.9851
27.8099
S2
f1 (ppm)

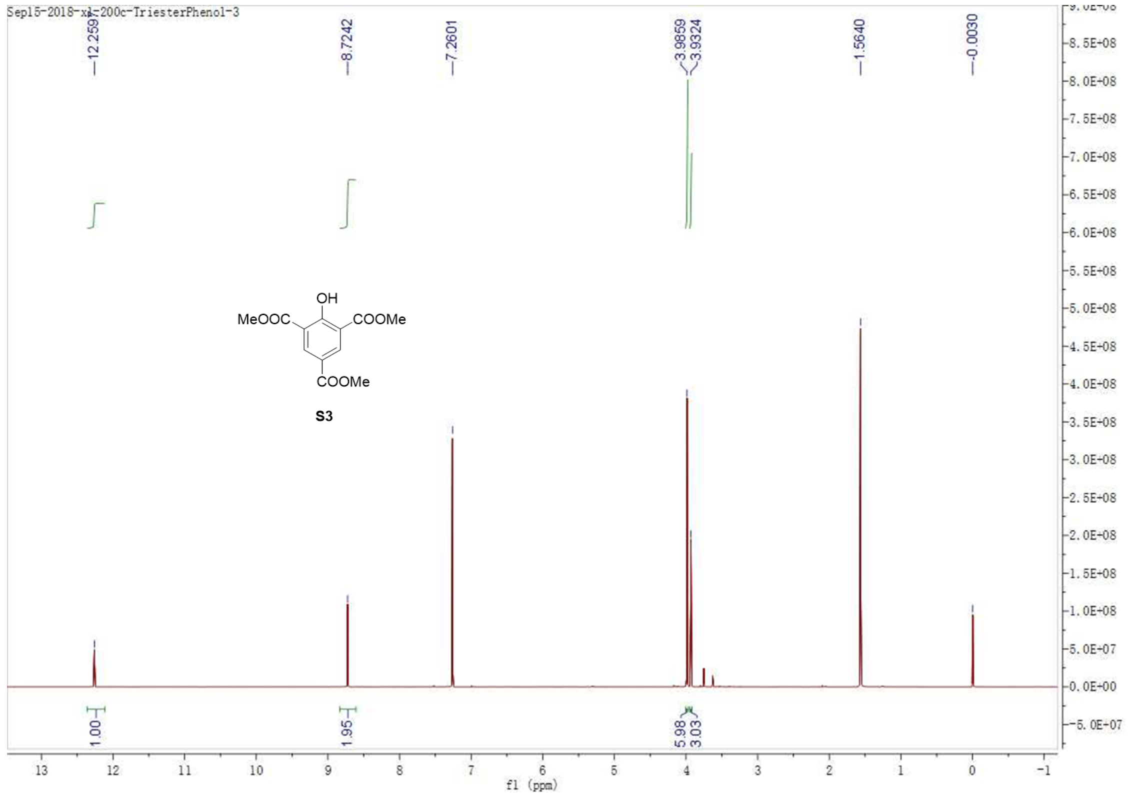
Sep15-2018-xl-200c-TriesterPhenol-3
12.2591
8.7242
7.2601
3.9859
3.9324
1.5640
-0.0030
1.00
1.95
5.98
3.03
f1 (ppm)
OH
MeOOC
COOMe
COOMe

S3

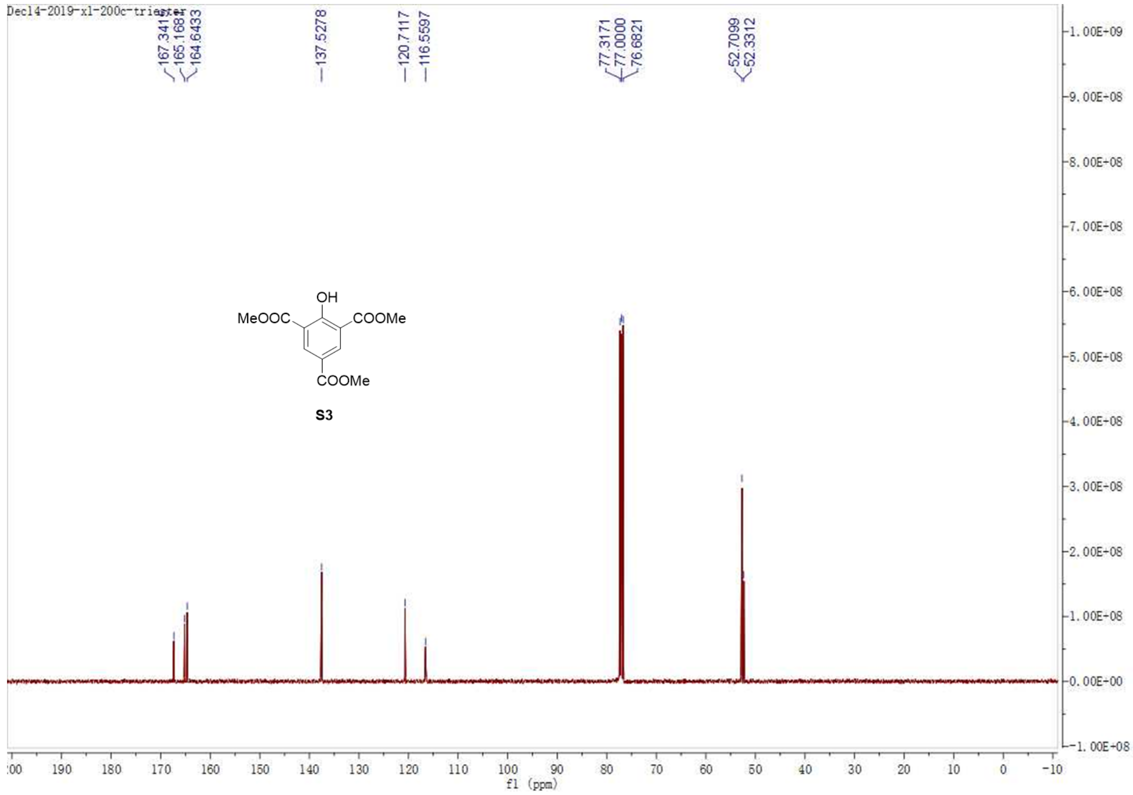
Dec14-2019-xl-200c-triester
167.3417
165.1681
164.6433
137.5278
120.7117
116.5597
77.3171
77.0000
76.6821
52.7099
52.3312
f1 (ppm)
OH
MeOOC
COOMe
COOMe

S3

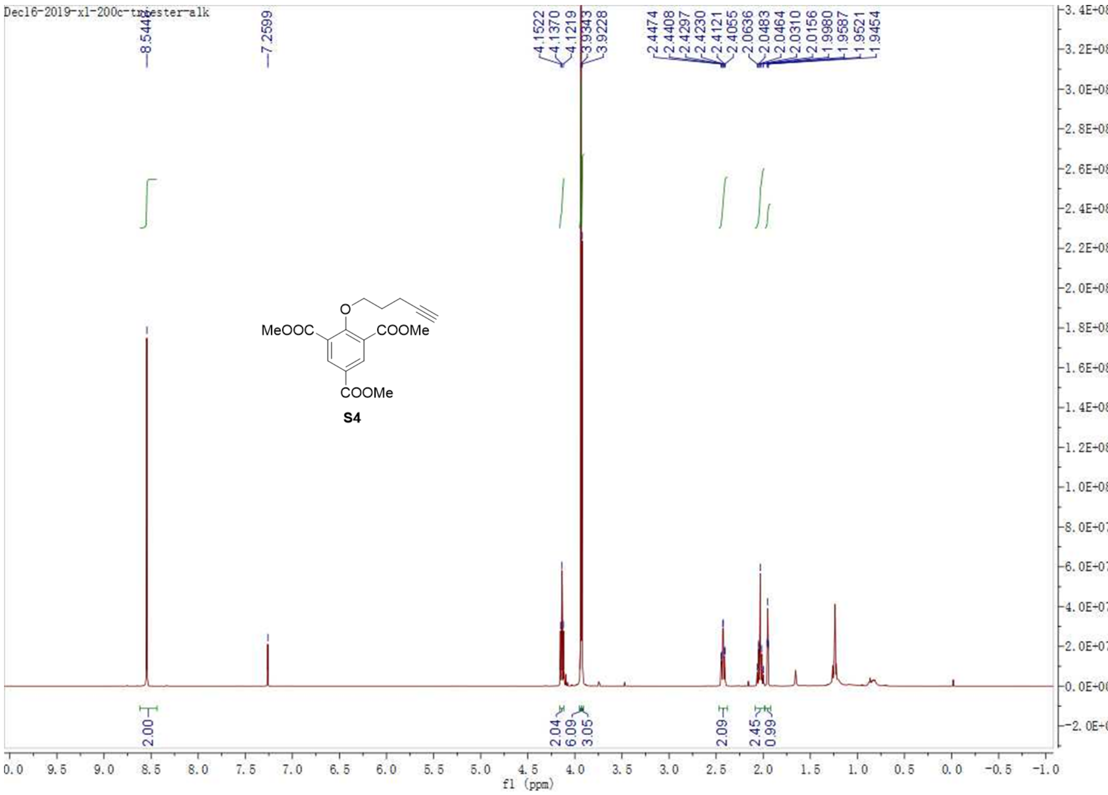
Dec16-2019-xl-200c-triester-alk
MeOOC
COOMe
COOMe
O
S4
f1 (ppm)

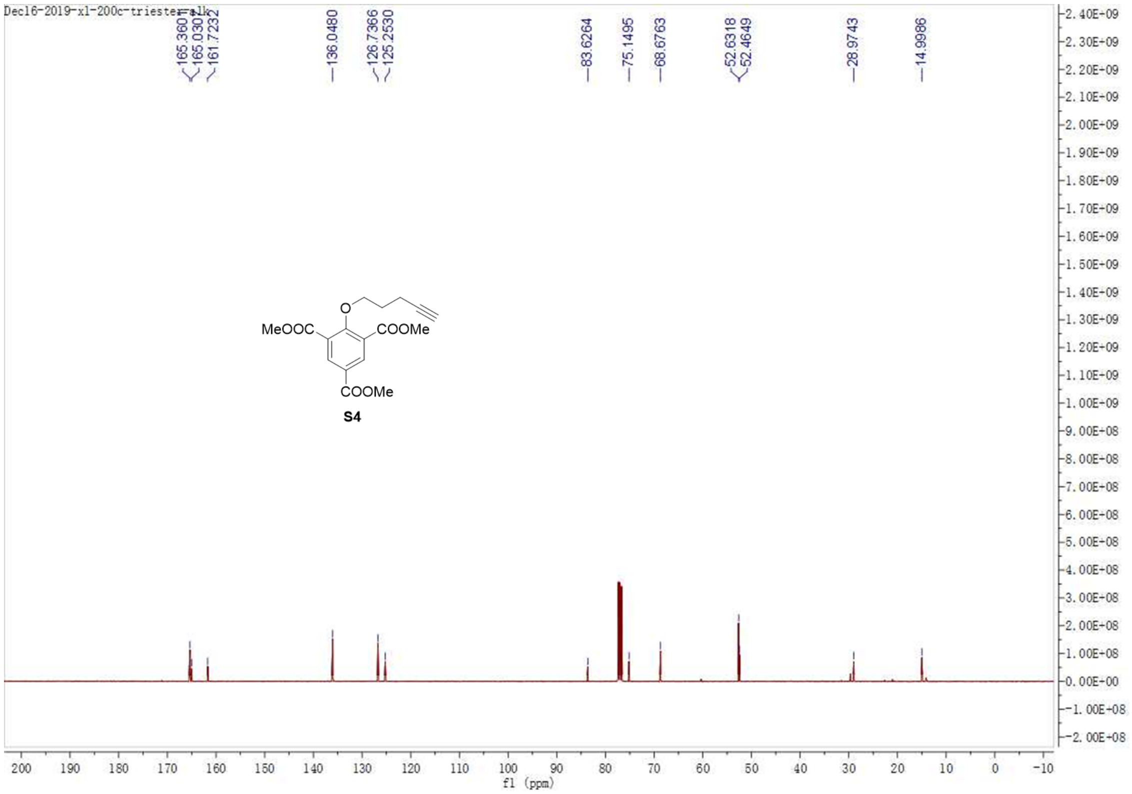
Dec16-2019-xl-200c-triester-alk
MeOOC
COOMe
COOMe
O
S4
f1 (ppm)

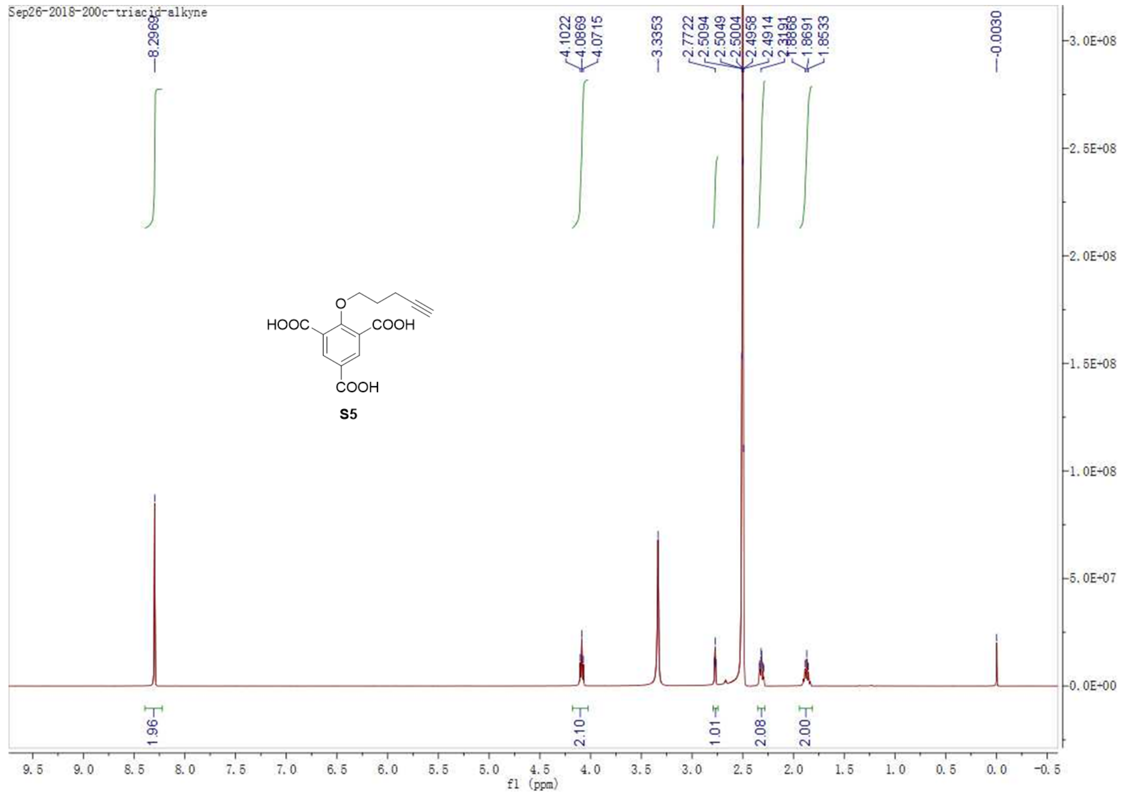
Sep26-2018-200c-triacid-alkyne
HOOC
COOH
COOH
S5
f1 (ppm)

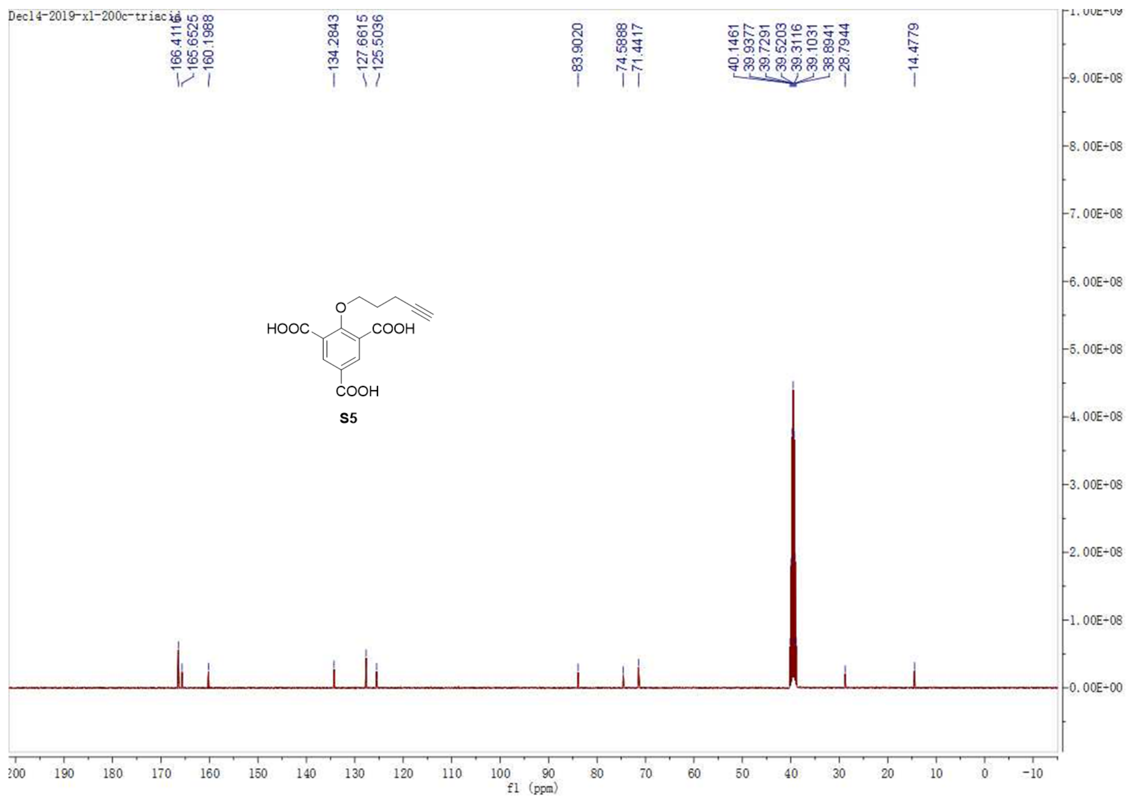
Dec14-2019-x1-200c-triacid
HOOC
COOH
COOH
S5
f1 (ppm)

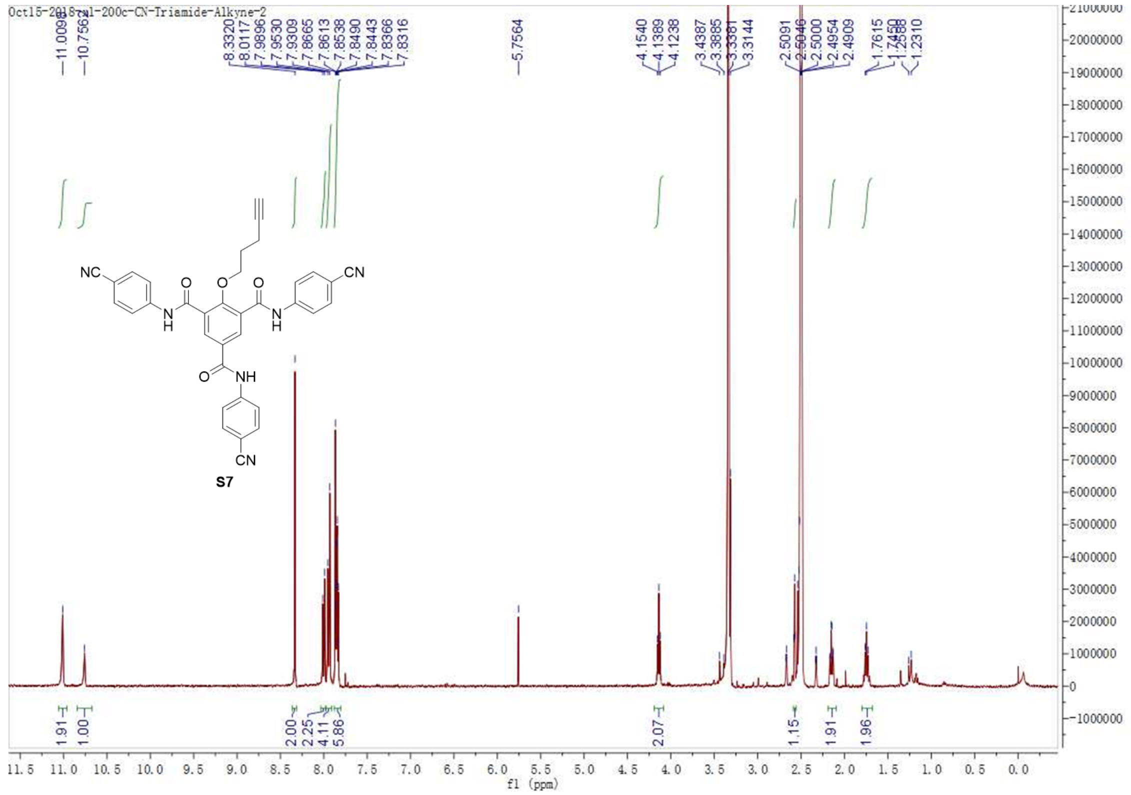

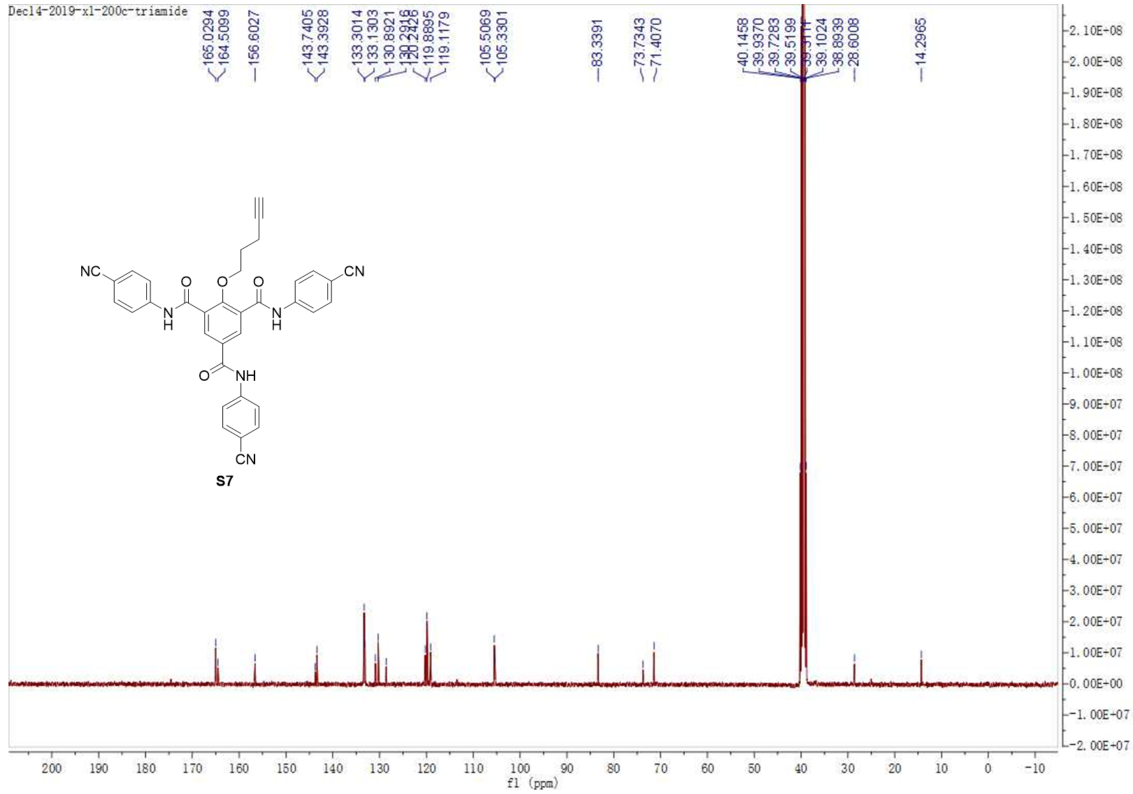

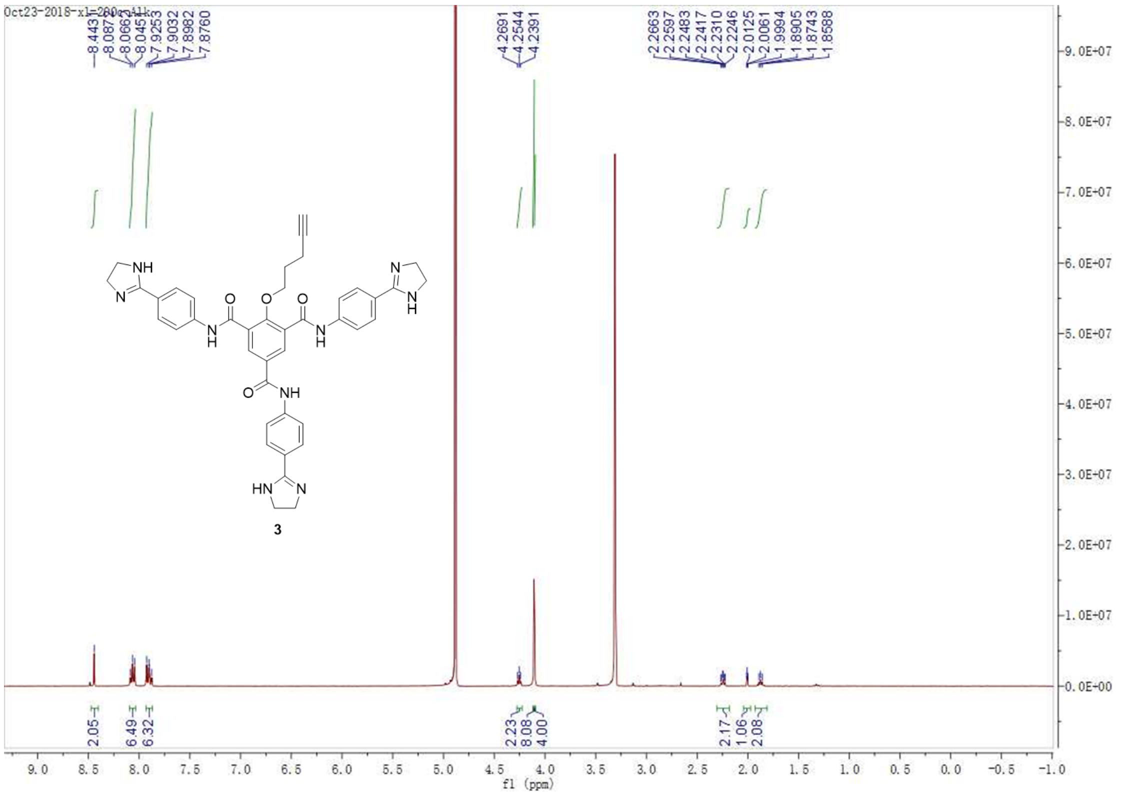
Oct23-2018-x1-200-alk
8.4431
8.0874
8.0662
8.0451
7.9253
7.9032
7.8982
7.8760
4.2691
4.2544
4.2391
2.2663
2.2597
2.2483
2.2417
2.2310
2.2246
2.0125
2.0061
1.9994
1.8905
1.8743
1.8588
2.05
6.49
6.32
2.23
8.08
4.00
2.17
1.06
2.08
NH
N
O
O
O
N
H
N
H
N
N
H
O
NH
HN
N
3
f1 (ppm)

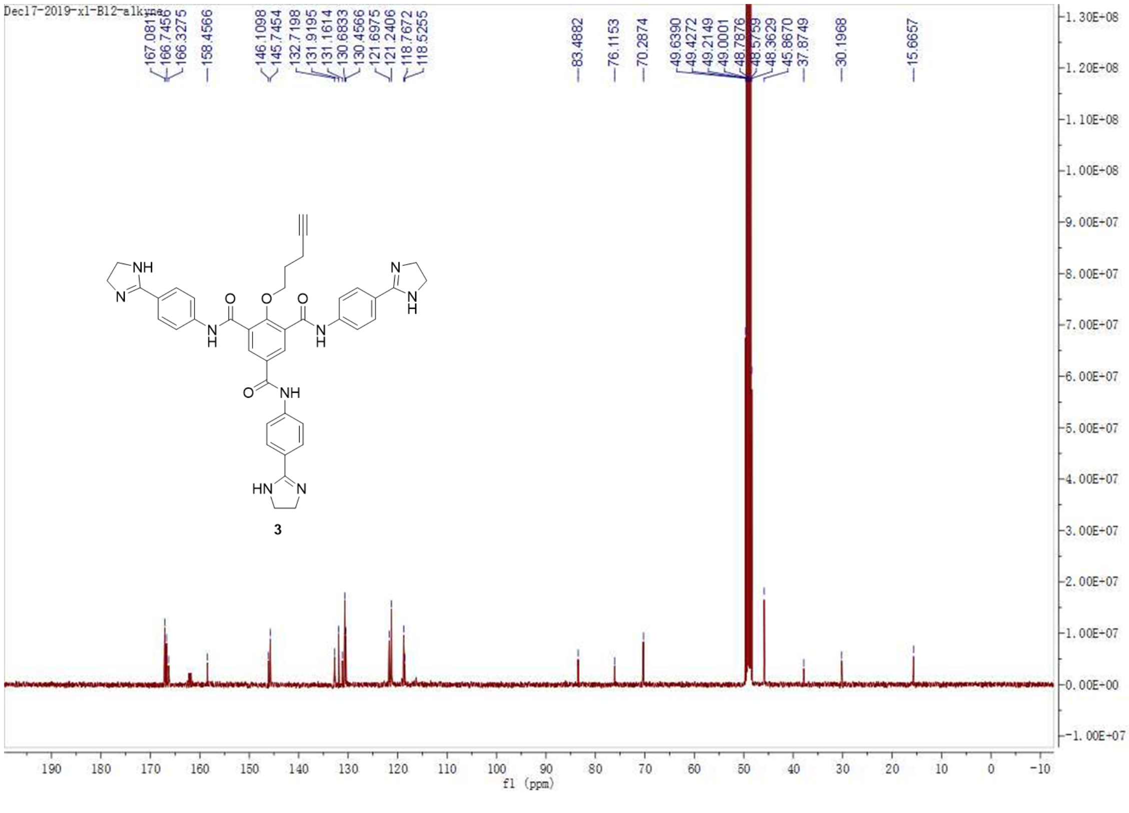
Dec17-2019-x1-B12-alkyne
167.0811
166.7450
166.3275
158.4566
146.1098
145.7454
132.7198
131.9195
131.1614
130.6833
130.4566
121.6975
121.2406
118.7672
118.5255
83.4882
76.1153
70.2874
49.6390
49.4272
49.2149
49.0001
48.7876
48.5759
48.3629
45.8670
37.8749
30.1968
15.6657
NH
N
O
O
O
N
H
N
H
N
N
H
O
NH
HN
N
3
f1 (ppm)

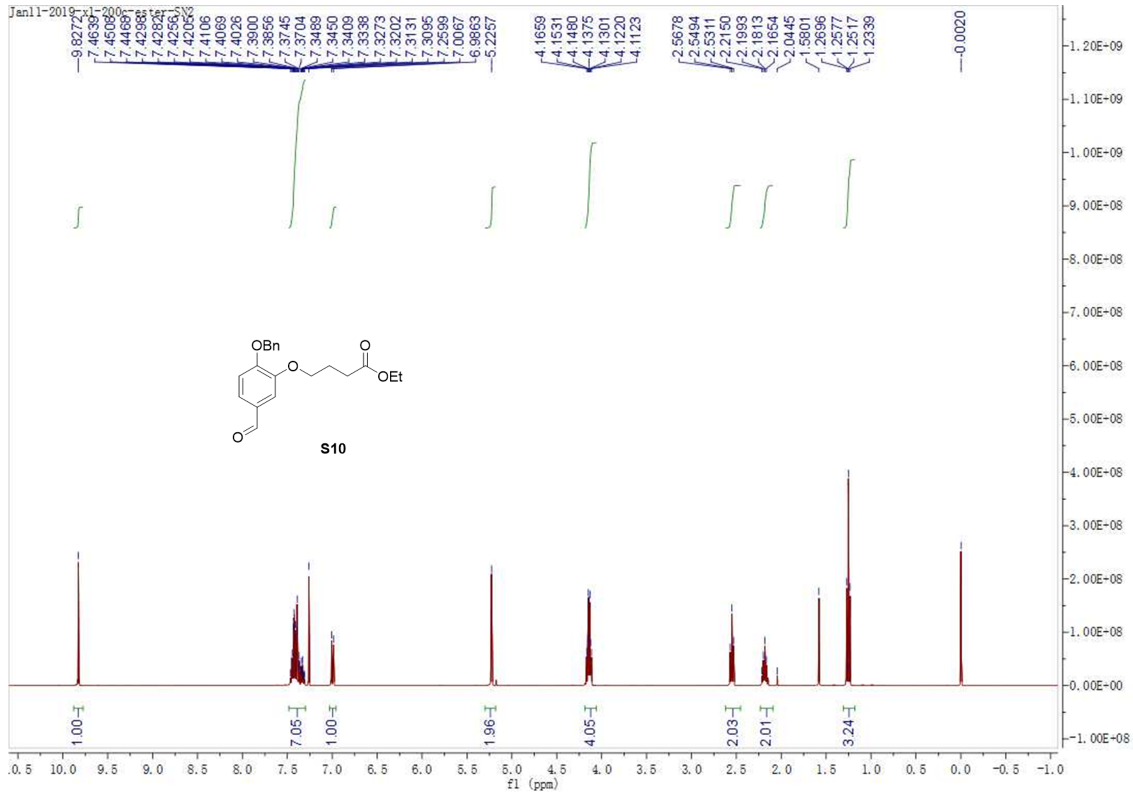

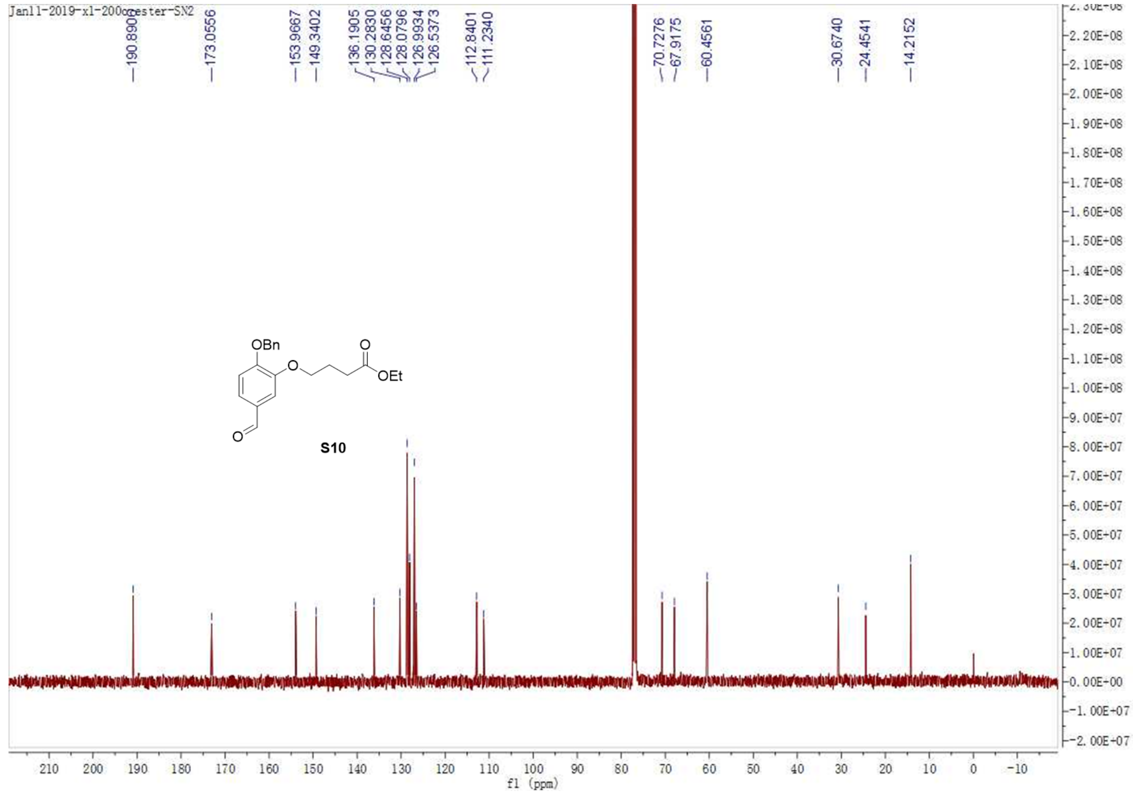

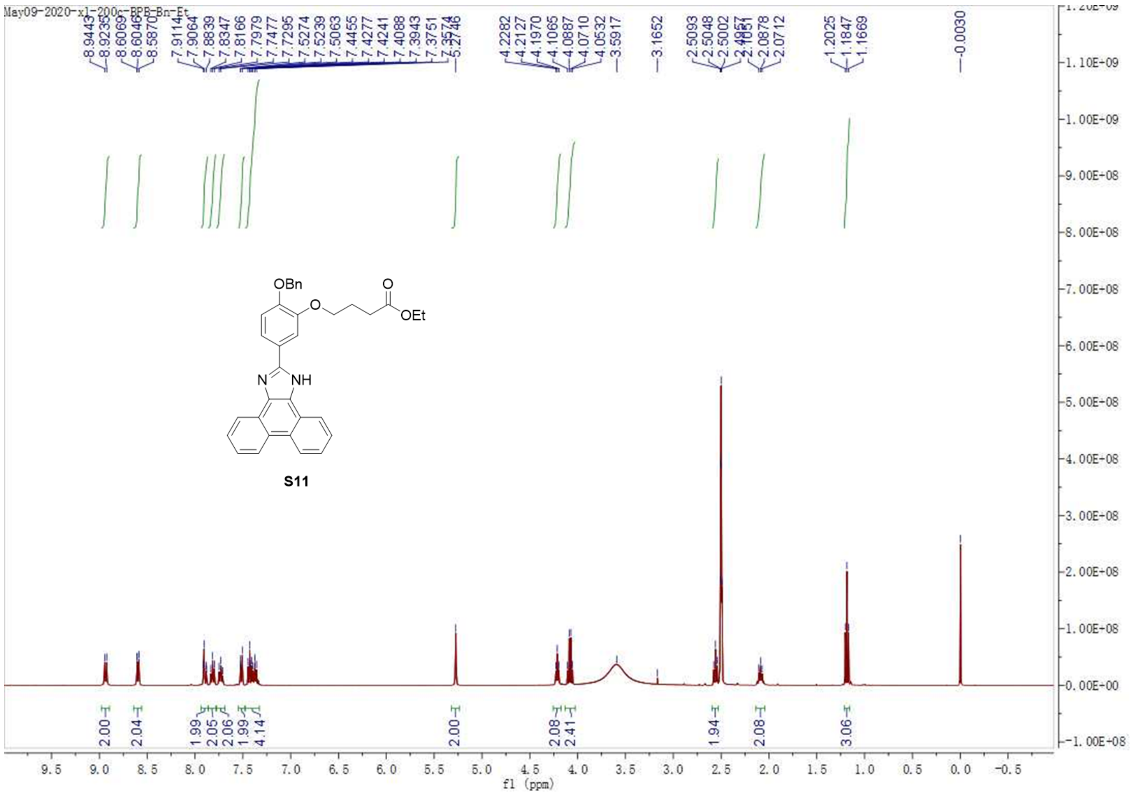

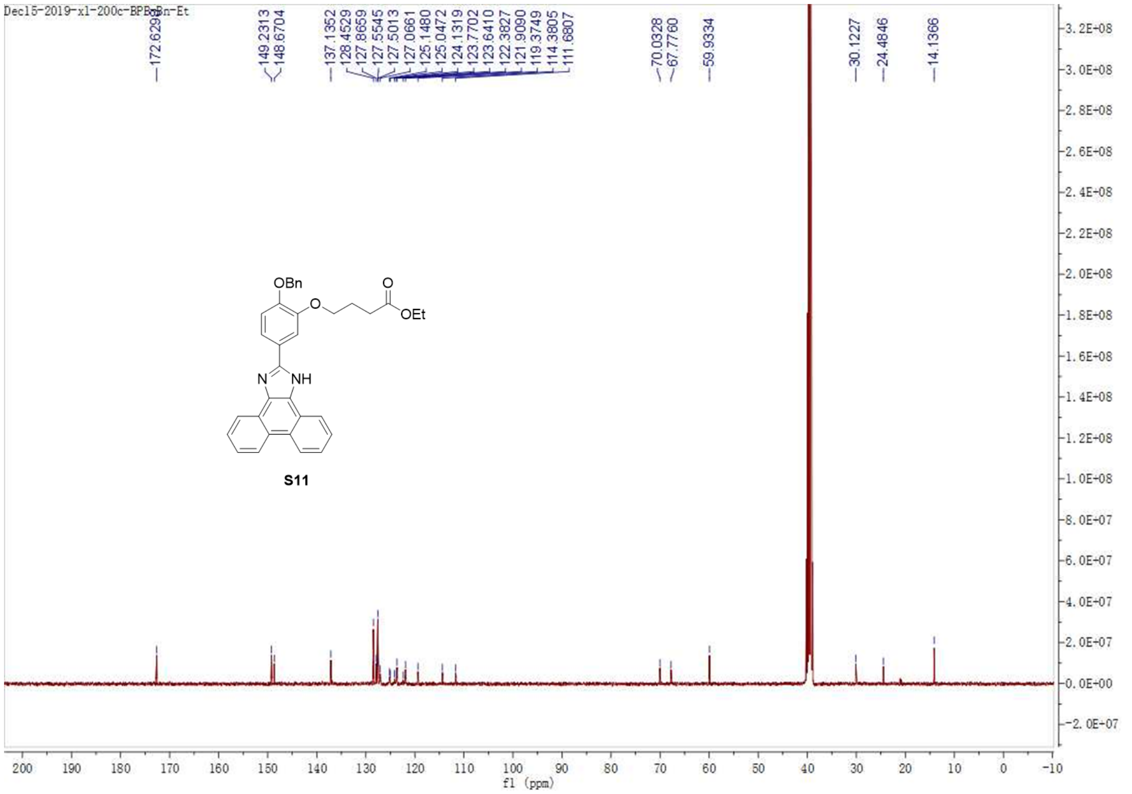

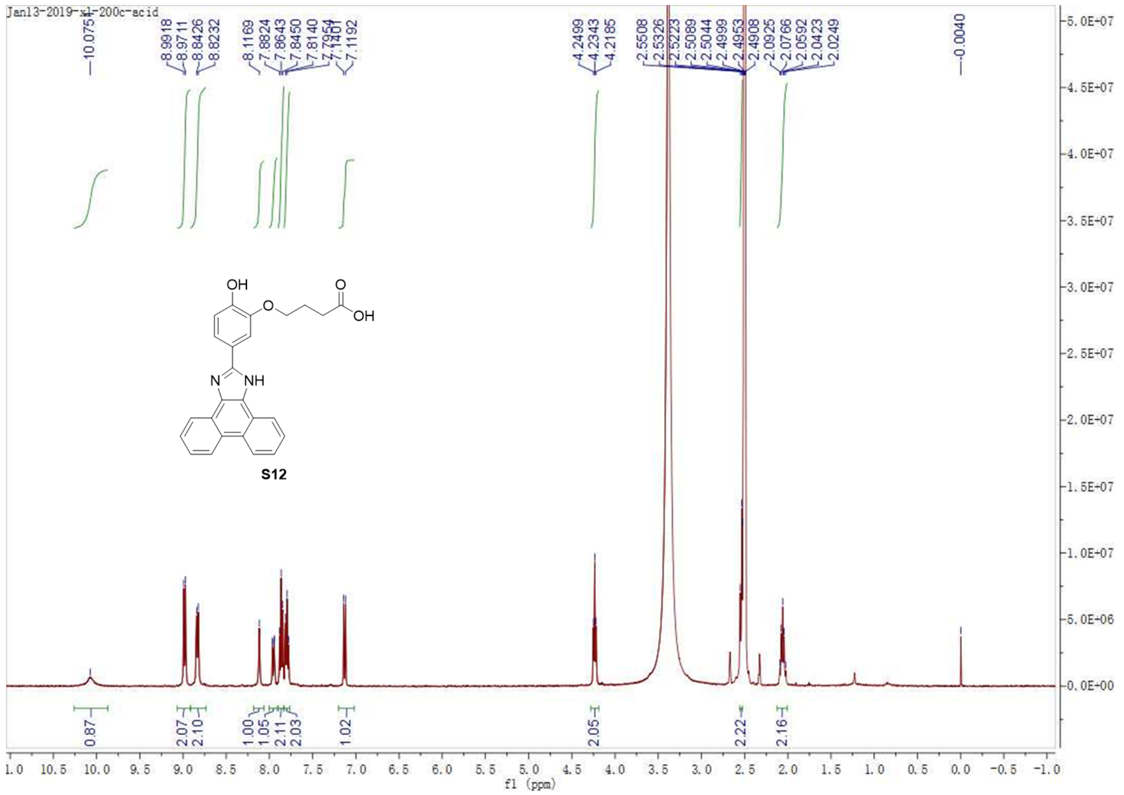
Jan13-2019-xl-200c-acid
10.0751
8.9918
8.9711
8.8426
8.8232
8.1169
7.8824
7.8643
7.8450
7.8140
7.7954
7.1401
7.1192
4.2499
4.2343
4.2185
2.5508
2.5326
2.5223
2.5089
2.5044
2.4999
2.4953
2.4908
2.0925
2.0766
2.0592
2.0423
2.0249
-0.0040
OH
O
OH
N
NH
S12
0.87
2.07
2.10
1.00
1.05
2.11
2.03
1.02
2.05
2.22
2.16
f1 (ppm)

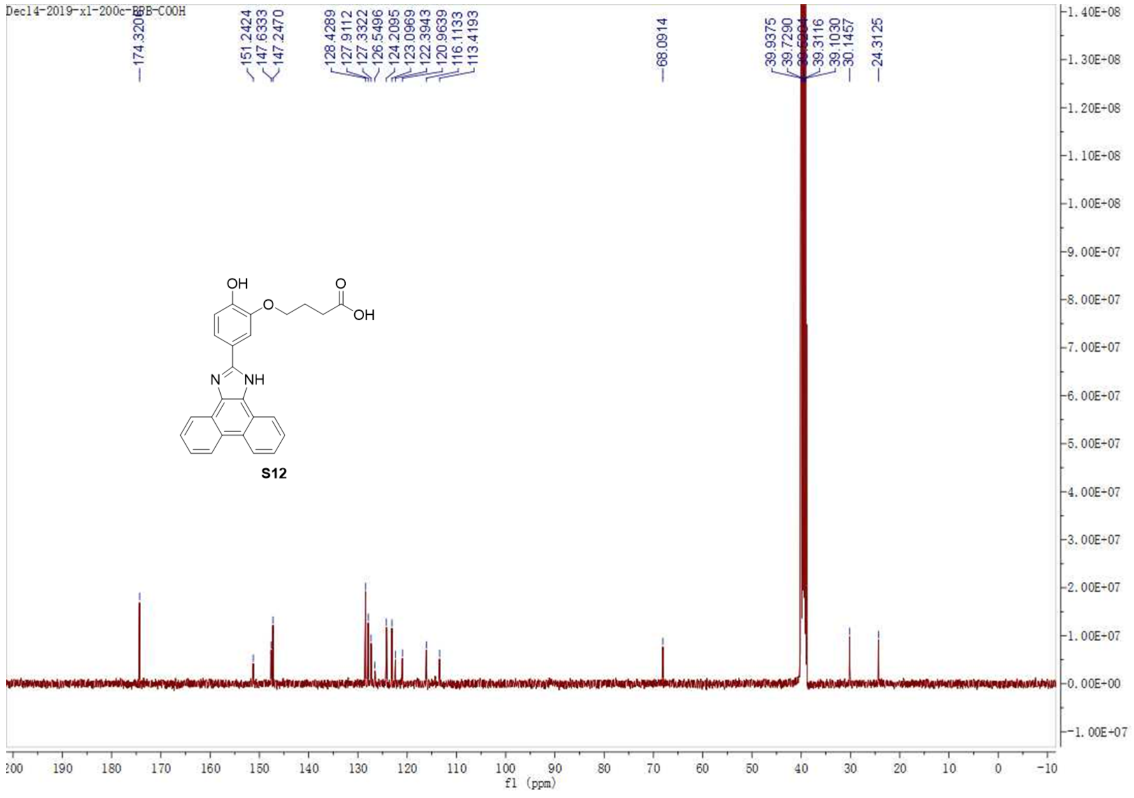
Dec14-2019-xl-200c-BPB-COOH
174.3206
151.2424
147.6333
147.2470
128.4289
127.9112
127.3322
126.5496
124.2095
123.0969
122.3943
120.9639
116.1133
113.4193
68.0914
39.9375
39.7290
39.3116
39.1030
30.1457
24.3125
OH
O
OH
N
NH
S12
f1 (ppm)
